# Interactions between emotion and action in the brain

**DOI:** 10.1101/812594

**Authors:** Liana C L Portugal, Rita C S Alves, Orlando Fernandes-Junior, Tiago A Sanchez, Izabela Mocaiber, Eliane Volchan, Fátima Erthal, Isabel A David, Jongwan Kim, Leticia Oliveira, Srikanth Padmala, Gang Chen, Luiz Pessoa, Mirtes G Pereira

**Affiliations:** Department of Physiology and Pharmacology, Laboratory of Neurophysiology of Behavior, Biomedical Institute, Federal Fluminense University, Niterói, RJ, Brazil; Laboratory of Neuroimaging and Psychophysiology, Department of Radiology, Faculty of Medicine, Federal University of Rio de Janeiro, Rio de Janeiro, RJ, Brazil; Laboratory of Cognitive Psychophysiology, Department of Natural Sciences, Institute of Humanities and Health, Federal Fluminense University, Rio das Ostras, RJ, Brazil; Laboratory of Neurobiology II, Institute of Biophysics Carlos Chagas Filho, Federal University of Rio de Janeiro, Rio de Janeiro, RJ, Brazil; Department of Psychology, University of Maryland, College Park, MD, USA; Scientific and Statistical Computing Core, National Institute of Mental Health, USA; Maryland Neuroimaging Center, University of Maryland, College Park, MD, USA

## Abstract

A growing literature supports the existence of interactions between emotion and action in the brain, and the central participation of the anterior midcingulate cortex (aMCC) in this regard. In the present functional magnetic resonance imaging study, we sought to investigate the role of self-relevance during such interactions by varying the context in which threating pictures were presented (with guns pointed towards or away from the observer). Participants performed a simple visual detection task following exposure to such stimuli. Except for voxelwise tests, we adopted a Bayesian analysis framework which evaluated evidence for the hypotheses of interest, given the data, in a continuous fashion. Behaviorally, our results demonstrated a valence by context interaction such that there was a tendency of speeding up responses to targets after viewing threat pictures directed towards the participant. In the brain, interaction patterns that paralleled those observed behaviorally were observed most notably in the middle temporal gyrus, supplementary motor area, precentral gyrus, and anterior insula. In these regions, activity was overall greater during threat conditions relative to neutral ones, and this effect was enhanced in the directed towards context. A valence by context interaction was observed in the aMCC too, where we also observed a correlation (across participants) of evoked responses and reaction time data. Taken together, our study revealed the context-sensitive engagement of motor-related areas during emotional perception, thus supporting the idea that emotion and action interact in important ways in the brain.

## Introduction

The processing of emotion-laden information, such as threat, is fast and prioritized. Emotion theories posit that emotions prime organisms for action tendencies (Darwin, 1872, Lang, Bradley & Cuthbert, 1997, Damasio, 1999) both in terms of approach and avoidance behaviors (Frijda, 1986). Darwin, for one, argued that emotions are adaptive insofar as they prompt actions that are beneficial to the organism. In line with this notion, manual approach-avoidance tasks with different types of apparatus, using a broad range of affective stimuli, showed that perceiving positive stimuli fosters approach behavior, whereas perceiving negative stimuli facilitates avoidance behavior (Chen and Bargh, 1999, Roelofs et al., 2009; Krieglmeyer & Deutsch, 2010; Saraiva et al., 2013, Phaf et al., 2014). Given the survival-related value of such actions, it is expected that emotional stimuli modulate signals in motor-related areas (Blakemore and Vuilleumier, 2017) much in the same way that they receive prioritized visual processing.

Further insight about emotion-motor interactions stems from studies utilizing transcranial magnetic stimulation and/or electromyography, which have reported increased excitability of the corticospinal tract during emotion perception (Oliveri et al., 2003, Hajcak et al., 2007, Coelho et al., 2010, van Loon et al., 2010, Nogueira-Campos et al., 2014). Furthermore, the emotional valence of a stimulus with which one is about to interact influences motor planning, as captured through the readiness potential, an electrophysiological marker of motor preparation (de Oliveira et al., 2012, Campagnoli et al., 2015). The existence of emotion-action interactions is also supported by functional neuroimaging studies of emotional modulation of motor-related brain areas (de Gelder et al., 2004, Grèzes, Pichon & de Gelder, 2007, Pichon, de Gelder & Grèzes, 2008, Pichon, de Gelder & Grèzes, 2009, Ahs et al., 2009, Pereira et al., 2010, Pichon, de Gelder & Grèzes, 2012, Kveraga et al., 2015, Kolesar, Kornelsen & Smith, 2017, Meyer, Padmala & Pessoa, 2019), even at the level of the spinal cord (Smith & Kornelsen, 2011, McIver, Kornelsen & Smith, 2013, Kornelsen, Smith & McIver, 2014).

More generally, the study of motor repertoires recruited by emotional stimuli has been explored in the non-human animal literature. For example, threat from predators or conspecifics prompts defensive behaviors including overt actions and/or immobility (e.g. Ratner, 1967; Blanchard and Blanchard, 1971). The corresponding human literature is relatively sparse, however. Characterization of defensive reactions in humans has capitalized upon studies of stabilometry, a methodology that assesses whole-body motor reactions (Azevedo et al., 2005; Bastos et al., 2016; Facchinetti et al., 2006; Volchan et al., 2011; for a review see Volchan et al., 2017). The findings from these studies point to similarities between non-human and human defensive behaviors, both of which exhibit motor defensive responses recruited according to threat imminence (Falselow and Lester, 1988). Particular features of threatening cues and associated context influence how threat imminence is perceived, determining the motor response to be triggered (Blanchard, Flannelly & Blanchard, 1986; Volchan et al., 2017). One such property is the self-relevance of the threat context, which can depend on subtle visual changes. For example, modifying the direction from a threat directed *away* to one directed *towards* the observer impacts threat perception. In fact, threat stimuli directed towards the observer are perceived as highly threatening, proximal, inescapable, and impossible to hide from (Fernandes et al., 2013, Bastos et al., 2016).

A brain area that is potentially a key site of emotion and motor interactions, and consequently involved in the selection of adaptive responses in different contexts, is the midcingulate cortex (MCC) (Pereira et al., 2010). One of the characteristics of the MCC is its prominent role in skeletomotor control (Vogt & Vogt, 2009), and the area appears to be especially involved with motor patterns that are context dependent (Talairach et al., 1973; Bancaud & Talairach, 1992). The MCC has two divisions, anterior (aMCC) and posterior (pMCC), with distinct functional profiles, cytoarchitecture and connections (for a review see Vogt, 2016). The aMCC has been described in the literature as an import site for cognitive control processes (Shackman et al., 2011), but also activated during fear- and pain-related conditions (Shackman et al., 2011; Vogt, 2016), while the pMCC exhibits almost no emotion-related activity (Vogt, 2005). In a previous study of our group (Pereira et al., 2010), we observed that the aMCC was recruited robustly only when participants performed a motor task in an unpleasant context, reinforcing the idea that the interplay between valence and motor information is important. In the study, aMCC activity during a target detection task paralleled the behavioral modulation associated with negative stimuli, such that increased activity in the negative context was sustained and correlated with the magnitude of the behavioral modulation.

Despite progress in understanding interactions between emotion and motor-related processing, the role of self-relevance during threat perception is poorly understood. In the present study, we addressed this issue using functional magnetic resonance imaging (fMRI) by adopting an experimental approach developed by our group in which behavioral and psychophysiological emotional responses varied as a function of stimulus relevance (Fernandes et al., 2013; Bastos et al 2016). In these studies, threatening visual stimuli consisted of pictures of a man holding a gun. Self-relevance was increased by changing gun direction from pointing away to pointing directly towards the participant. Here, we investigated brain responses while participants performed a task following exposure to such stimuli. We hypothesized that viewing threat stimuli directed towards the observer would recruit emotional and motor areas more robustly. In particular, some of these areas would exhibit a valence by context interaction, such that the differential response to threat would be enhanced when stimuli were self-relevant.

## Methods

### Participants

Forty-nine right-handed undergraduate or graduate students participated in the study (26 females, mean age=26.6, standard deviation=5.0). Six participants were excluded from analyses due to excessive errors (>25%, mean error rate was 5.5%) leading to a final sample of 43 subjects for reaction time data analysis (20 females, mean age=26.7, standard deviation=4.8). Additionally, two volunteers were excluded due to data loss during data transfer, two due to excessive head movement during scanning (greater than 6 mm) and one because of poor structural-functional alignment, leading to a final sample of 38 participants (17 females, mean age=27.1, standard deviation=4.8). All participants had normal or corrected-to-normal vision, reported no psychiatric or neurologic problems and were not taking medication with central nervous system action. The project was approved by the local ethics committee of Federal Fluminense University, Rio de Janeiro, Brazil, and each participant gave written informed consent prior to participation.

In a previous study, we reported findings using 26 participants of the sample reported here (Fernandes et al., 2017). However, the study focused on an entirely distinct question, namely it employed machine learning to predict individual-level negative-affect trait measures from brain activation.

### Stimuli

Eighty-four pictures were employed. Pictures were either obtained from the World Wide Web, purchased from Getty Images® (http://www.gettyimages.com), or produced by the authors with support of a professional photographer, except for one picture that was obtained from the International Affective Picture System (IAPS) (Lang, Bradley & Cuthbert, 2005). The stimulus size was standardized to 1024 × 768 pixels.

The 84 pictures consisted of two sets: threat and neutral stimuli (42 pictures each). Threat stimuli were pictures of a man holding a gun. Neutral stimuli were pictures of a man holding an object, such as a camera or a domestic tool. In both sets, the guns or the neutral objects were either directed towards or away from the participant (21 pictures in each). In all, there were a total of four picture categories: (1) threat stimuli directed towards the participant; (2) threat stimuli directed away from the participant; (3) neutral stimuli directed towards the participant; and (4) neutral stimuli directed away from the participant. The pictures were matched along several properties to minimize potentially confounding effects (Steinmetz et al., 2011). The ethnicity of men holding guns or non-lethal objects was balanced. Threat and neutral stimuli were also matched in terms of physical properties such as brightness, contrast, and spatial frequency (Table S1 and S2, supplemental material). Electrodermal responses and evaluative characterization of stimuli are depicted in supplemental material, and revealed that our threat manipulation was successful: threat directed towards the participant was judged as more intense, nearer and inescapable, and providing reduced possibility of hiding. Additionally, threat stimuli directed towards the participant induced greater skin conductance response compared to all conditions, indicating an increased recruitment of sympathetic system in this context.

### Experimental design and procedure

Stimuli were back-projected onto a screen located in front of the participant’s body and were viewed inside the scanner using a mirror attached to the head coil. The stimuli were presented using Presentation software (Neurobehavioral Systems, version 11.0, Inc., Albany, CA, USA). The experimental session was divided into four runs. Each run consisted of 14 blocks. The order of blocks of each category (threat stimuli directed towards or away from the participant, and neutral stimuli directed towards or away from the participant) was randomized. Each block consisted of three different pictures (5 seconds each) of the same category presented in sequence, followed by a 12-second fixation cross. Each trial began with the presentation of the picture (740×520 mm) together with a central fixation cross (9×9 mm). Three seconds after picture onset, a square (cue, 35×35 mm) appeared around the fixation cross, indicating that the target would appear at any moment. The target, a small central circle (inner circumference diameter of approximately 28 mm), appeared 700-1,200 ms after the square, and both remained on until the end of the trial. The fixation cross, the cue, and target were shown over the picture. Participants were instructed to attend to each picture while maintaining their eyes at a fixation spot at the centre of the screen, and to press a button with their right index finger as quickly as possible following target onset. An MR-compatible response key, positioned on the right side of the participant’s abdomen, recorded the responses. Each trial during which one picture was shown followed by target detection lasted for 5 s (Fig. 1). Each run was 378 seconds long.

**Fig. 1.**
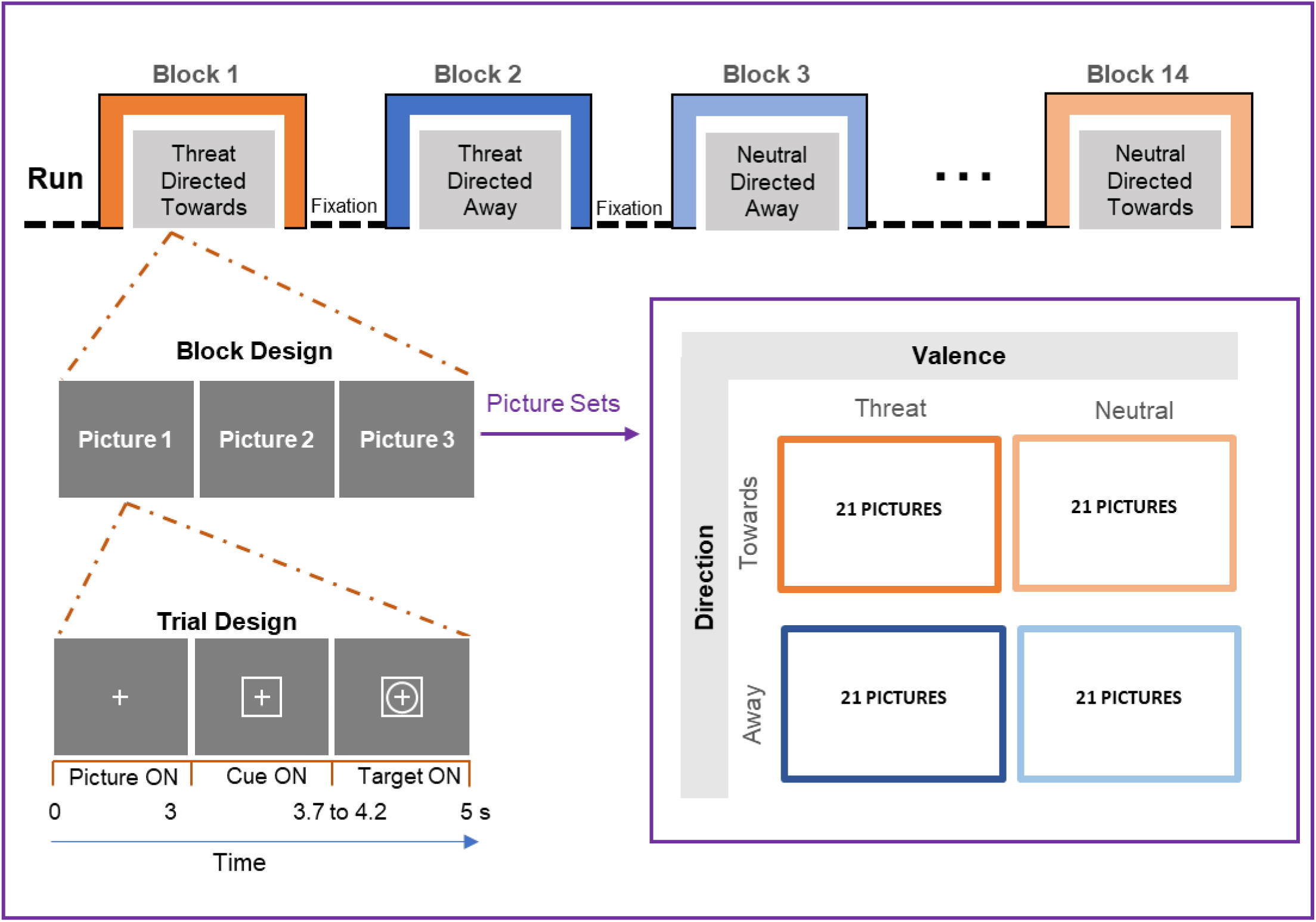
Experimental design. Each block contained three trials, each of which began with the presentation of a picture together with a central fixation cross. Three seconds after picture onset, a square (cue) appeared around the fixation cross, indicating that the target would appear at any moment. The target, a small central circle, appeared 700-1,200 ms after the square, and both remained on until the end of the trial. The fixation cross, the cue, and target were shown over the picture. Twenty one threat stimuli directed towards and directed away from the participant, and neutral stimuli directed towards and directed away from the participant were presented. Threat stimuli were pictures of a man holding a gun. Neutral stimuli were pictures of a man holding an object, such as a camera or a domestic tool.

At the beginning of the session, during anatomical scanning, the participants performed a practice task, which was similar to the main experiment except that all images involved pictures of neutral objects, such as tools and furniture, and feedback was displayed on the screen.

### Image acquisition

Functional and anatomical MRI data were collected at the Department of Radiology at Hospital Universitário Clementino Fraga Filho (Federal University of Rio de Janeiro) on a 1.5T Siemens (Magnetom Avanto) scanner. The fMRI runs were acquired on a sequential ascending order, using a gradient echo EPI single-shot sequence covering 25 axial slices (4-mm-thick; 0.6-mm gap; TR/TE=2000/40 ms; IST=80 ms; FOV=256 mm; matrix, 64×64; voxel dimensions, 4×4×4.6 mm). Head movements were restrained by foam padding. In each run, 198 functional volumes were acquired in a total of four runs. In addition, a three-dimensional high-resolution T1-weighted anatomical image (TR/TE=2730/3.27 ms; 128 slices; 0.6-mm gap; FOV=250 mm; voxel dimensions 1.33 × 1 × 1.33 mm) was obtained at the beginning of the session for functional-to-anatomical image registration.

### Functional MRI preprocessing

Preprocessing of the functional and anatomical MRI data used AFNI (Cox, 1996; http://afni.nimh.nih.gov/) and the Statistical Parametric Mapping software package (SPM8, Friston et al., 1995; Wellcome Department of Cognitive Neurology, London, UK). The first three volumes of each functional run were discarded to account for equilibration effects. Slice-timing correction used Fourier interpolation (AFNI 3dTshift) to align the onset times of every slice in a volume to the first slice. A six-parameter rigid body transformation (AFNI 3dvolvreg) was used to correct head motion within and between runs by spatially registering each volume to the first volume.

The SPM8 package was used to skull strip the high-resolution anatomical images. Anatomical images were rotated to match the oblique plane of the functional data using AFNI 3dWarp. Each participant’s anatomical scan was registered to the TT_N27 template of the AFNI package for normalization to Talairach space (Talairach and Tournoux, 1988). The same transformation was applied to the functional data. An 8-mm full-width half-maximum Gaussian filter was used to spatially smooth all volumes, and the average intensity at each voxel (for each run) was scaled to 100.

### fMRI analysis

Data analysis was performed according to multiple linear regression as implemented in AFNI (using 3dDeconvolve). The first (fixed) level involved determining the regression coefficients of variables of interest, which modelled the effects of each experimental condition: threat stimuli directed towards, threat stimuli directed away, neutral stimuli directed towards, and neutral stimuli directed away. Before estimation via multiple regression, regressors of interest were convolved with a canonical haemodynamic response function (GAM in AFNI; Cohen, 1997). The 2 seconds following the cue onset were used as regressors of interest for each condition, and the 12-second fixation cross between blocks was considered as the baseline. The four runs were modelled together. Movement parameters, and their first derivatives, were entered as covariates of no interest associated with the participants’ head motion. To further control for head motion, we excluded volumes with a frame-to-frame displacement of more than half of the voxel size from the analysis (Siegel et al, 2014). Only one participant had four volumes excluded.

Second-level group analyses were conducted by means of repeated-measures ANOVAs with the factors of valence (threat, neutral) and context (directed towards, directed away). As our central aim was to investigate valence by context interactions, we first identified brain areas modulated by valence (note that this main effect is orthogonal to the interaction term). To do so, the main effect of valence was employed at a voxel-level threshold of 0.01 (corrected based on False Discovery Rate). These areas were considered as regions of interest (ROIs) for subsequent analysis, thereby reducing the multiple comparisons problem. Mean regression coefficients for all significant voxels (cluster of activation for the main effect of valence) were extracted for each subject and condition. Each ROI was further interrogated for the valence by context interaction effect. In addition, brain regions with a valence by context interaction effect that survived correction at a voxel-level threshold of 0.05 are reported.

Given the potential role of the aMCC for the integration of emotional and motor signals, activity in this region was further investigated using an anatomical atlas-based ROI. The aMCC ROI was created based on the Destrieux et al. (2010) atlas, which subdivides the cingulum into several segments following the antero-posterior direction, as proposed by Vogt (Vogt, Berger & Derbyshare, 2003, Vogt, Vogt & Laureys, 2006). Mean regression coefficient values, for each subject and condition, were extracted and tested for evidence of a valence by context interaction effect (no thresholding based on functional data was applied).

### Bayesian statistical analysis

The null hypothesis significance testing (NHST) framework has come under increased scrutiny in recent years. In particular, the hard threshold of 0.05 has come under attack, with reasonable researchers calling for stricter thresholds (Benjamin et al., 2018) or, conversely, for the dichotomous use of p-values to be abandoned (McShane et al., 2017). We do not consider a binary threshold to be satisfactory and believe that p-values should be treated continuously. In fact, in the present paper, wherever possible, we employed Bayesian statistical analysis (i.e, all analyses except the voxelwise one described above).

As Bayesian analysis is not as widely used, we briefly compare this framework to NHST. Consider a scenario in which a single one-sample *t*-test with 20 degrees of freedom (e.g., 21 subjects) is employed in the NHST setting. The null hypothesis H_0_ is that the population mean is zero. Suppose that the data indicate that t_20_ = 2.85. If H_0_ were true, the probability of observing a t_20_-value so large is rather low (0.01 in Fig. 2, left). The t_20_-value thus provides a measure of “surprise”: How surprising would it be to observe such an extreme value in a world in which H_0_ were really true? The extent of surprise corresponds, of course, to P(data | H_0_). By custom if P < 0.05, then one declares that the effect is “statistically significant”.

**Fig. 2.**
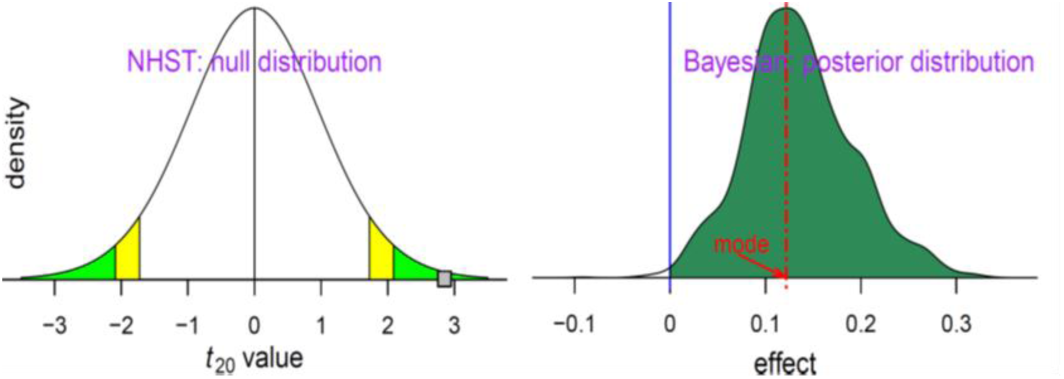
Different probability definitions and goals of conventional and Bayesian frameworks. Left: Statistical inferences under null hypothesis significance testing are based on how extreme the data are in the context of the null hypothesis. Green/yellow tails symbolize a two-sided significance level of 0.05/0.1. In the example, the data produces a t-value of 2.85 (small gray square). Right: Inferences under the Bayesian framework directly address the question of research interest: what is the probability of the effect magnitude being greater than 0 with the data at hand? The “output” in a Bayesian inference comprises the entire posterior density function, which summarizes the uncertainty in estimating the effect magnitude.

The Bayesian framework aims to answer a different, though, related question: what is the probability of a research hypothesis H given the data, P(H | data)? Note what is being “measured” and what is the “given” in this proposition, as opposed to the preceding NHST formulation (i.e., P(data | H_0_)). Such a probability can be computed by using Bayes’ rule (for an introductory text, see Kruschke, 2010). In a typical setting, the research hypothesis is a statement that refers to an effect or parameter being estimated (e.g., mean, difference of means, etc.). An attractive property of this framework is that it is not typically formulated to generate a binary decision (“real effect” vs “noise”, or “significant” vs. “not significant”) but instead to obtain the entire probability density distribution associated with P(θ | data), where θ is the parameter being estimated (Fig. 2, right). This so-called posterior distribution is interpreted in a natural way, although it may take getting used to for those who are unfamiliar with Bayesian inferences. For example, P(θ > 0 | data), which we call P+, is the area under the curve in Fig. 2 (right); in the present case, P+ = 0.99, namely, there is very strong evidence that the effect of interest (e.g., mean, difference of means, etc.) is greater than zero (conditional on the data, the prior distribution, and the model). Note that small values of P+ convey support for a negative effect; for example, P+ = 0.01 indicates that the probability of the effect being positive is only 0.01, which implies that the probability of it being negative is 0.99. Finally, the estimation of posterior distributions requires the specification of a prior distribution. In all analyses reported here, we employed so-called “uninformative” priors, which accordingly do not have a notable impact on the conclusions.

The posterior distribution provides a summary description of the likelihood of observing parameter values given the data, so it naturally conveys variability. Some authors use cut-off points to summarize “strong”, “moderate”, or “weak” evidence, but we encourage an approach that both quantifies and qualifies the evidence, without making decisions in terms of “passes threshold” versus “fails to pass threshold”. Note that we do not employ Bayes factors, which some have advocated as a potential feature of Bayesian modeling. Because Bayes factors consider the probability of “null” effects (e.g., a mean of zero) versus an “alternative” effect (e.g., a mean different from zero), we believe it is also problematic, because formulating the problem in terms of “null” effects of exactly zero is often unrealistic (because effects of experimental manipulations are seldom zero), thus largely inflating the evidence for the alternative hypothesis (thus creating “large” Bayes factors). See Chen et al. (2019a) for further discussion. Finally, given that we do not view thresholding as adequate, the P+ probability values that we provide are (by definition) “one-sided”. For readers who absolutely insist on comparing P+ values with standard cut-offs that are “two-sided”, they should bear in mind our definition.

### Bayesian analysis of reaction time and individual ROIs

Bayesian analyses were performed via the brms R package (Bürkner, 2017), which employs the Stan probabilistic language to compute posterior distributions via state-of-the-art Monte Carlo Markov Chain modelling (Stan Development Team, 2016). To analyze the effects of valence, context, and their interaction, we ran three separate Bayesian analyses: the two main effects and the interaction. If we consider the four levels of the experimental variables (in order: threat_towards, neutral_towards, threat_away, neutral_away), the effect of valence can be obtained via the contrast expression (+1 –1 +1 –1), the effect of context via the contrast expression (+1 +1 –1 –1), and their interaction via the contrast (+1 –1 –1 +1). Given this coding, the three models were simply of the form effect ∼ Normal(α, σ) σ ∼ Normal(0,10) where “∼” indicates “distributed as”, α is a single intercept, σ is the residual variation, and the last line specifies the prior distribution.

### Region-based Bayesian analysis

A standard ROI analysis evaluates the effect of interest for each ROI, separately. Given the multiplicity of ROIs, investigators commonly perform some correction for multiple comparisons, say via Bonferroni correction. Here, we analyzed the interaction between valence and context by performing a Bayesian multilevel analysis of the ROI data by using the Region-Based Analysis (RBA) program of the AFNI suite (Chen et al., 2019b). In this approach, the data from all ROIs are included in a single multilevel model that evaluates the effects of interest. By doing so, the contributions to fMRI signals of subject-level effects (i.e., subject effect across conditions), and ROI-level effects (i.e., ROI effect across subjects), can be accounted for in a model that simultaneously ascertains the interaction effect.

Thus, under the standard general linear model, the effect at each region is estimated independently from other regions; there is no information shared across regions, hence, the multiple comparisons step. In contrast, in the Bayesian multilevel framework the effects across regions are shared (technically, via partial pooling) by assuming that the ROI contributions are normally distributed. The latter implies a Gaussian prior distribution on the ROI effects. The “output” of the Bayesian multilevel model comprises only one overall posterior that is formulated as a joint distribution in a high-dimensional parameter space (thus, no “correction” is needed). For summary purposes, posteriors of the effects for every ROI can be plotted separately; but they are not independent and technically are simply marginal distributions (that is, projections along particular variables). For formal details of the approach adopted here, please refer to Chen et al. (2019b); for a less technical exposition, see Chen et al. (2019a).

### Brain-behavior correlation

To test whether the magnitude of the emotional modulation in the aMCC (anatomical ROI) was related to the magnitude of the behavioral emotional modulation, we used a modulation index for both measures. In the case of behavior, the index was defined as the mean RT to targets during threat trials minus the mean RT to targets during neutral trials, for the directed towards and directed away contexts, separately. Accordingly, negative values of the index indicated that participants were faster to detect the target when threat stimuli were presented; positive values indicated the reverse. In the case of aMCC responses, we computed an analogous index based on the average regression coefficient values for each condition of interest.

We performed Bayesian analysis of the brain-behavior relationship for the right and left aMCC, separately, by using the brms R/Stan package. Because this involved a linear relationship, the model was simply

aMCC ∼ Normal(μ, σ)

μ ∼ α + βxRT

α ∼ Normal(0,10)

β ∼ Normal(0,10)

σ ∼ HalfCauchy(10)

where the second line specifies the linear association, and the last three indicate the prior distributions.

## Results

The analyses reported next were all performed within a Bayesian framework, except the voxelwise analyses which followed the standard statistical approach employed in neuroimaging. Here, we report the extent of evidence, P(θ > 0 | data), as P+. Values of P+ closer to 1 indicate stronger evidence that the effect of interest (e.g., mean, difference of means, etc.) is greater than zero (see Methods). Small values of P+ convey the extent of support that the effect is negative; that is, the probability that the effect is negative is given by (1 – P+).

### Behavioural performance

Behaviourally, we observed a tendency for the speeding up of RT to targets when viewing negative pictures (467 ms) relative to those following neutral pictures (474 ms) during directed towards trials (effect size d = -0.39). During directed away trials, mean RTs were the same (474 ms) for threat and neutral trials (effect size d= - 0.02). The Bayesian analysis of RTs revealed evidence for a valence by context interaction (P+ = 0.040), an effect of valence (P+ = 0.038), and an effect of context (P+ = 0.051). Figure 3 shows the RT data together with the Bayesian posterior density plots. In all three cases, because most of the evidence for the effect of interest is negative, the P+ values are close to zero.

**Fig. 3.**
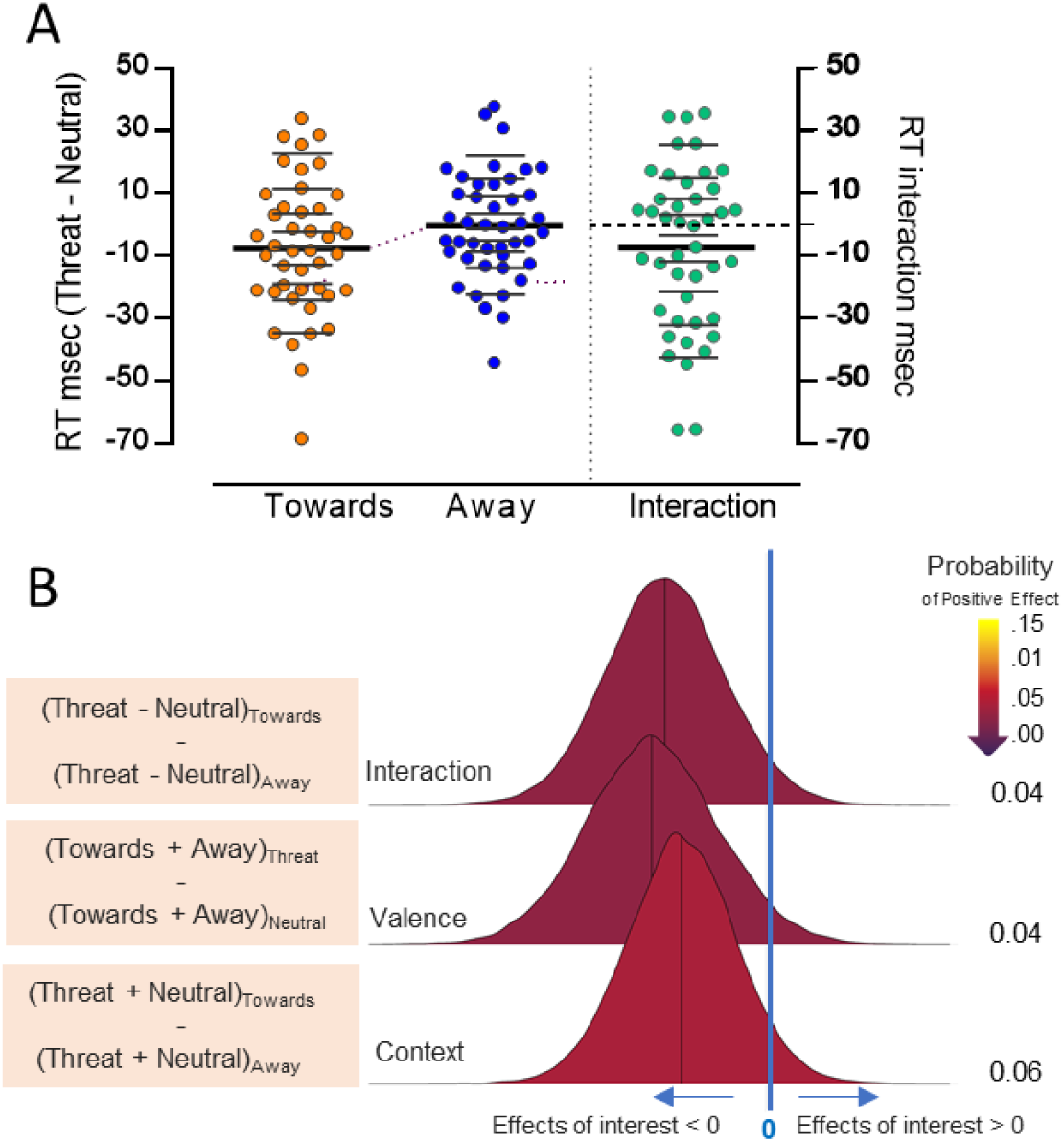
Behavioral data and Bayesian posterior density plots. (A) The circles show reaction time (RT, ms) differences (threat minus neutral), and a behavioral interaction index (RT difference during directed towards minus RT difference during directed away). Thicker lines mark the mean and thinner lines the deciles for each experimental condition. The thin purple line marks the shift between means. Data illustrations used some of the graphical tools proposed by Rousselet et al (Rousselet, Pernet & Wilcox, 2017). (B) Bayesian posterior density plots of the valence by context interaction effect, the main effect of valence, and the main effect of context for RT data. Inferences under the Bayesian framework directly address the probability of the effect magnitude being greater than 0, which we call P+. The color bar represents P+. Note that the reddish to dark purple color associated with small values of P+ convey support that the effect is negative; in this case, P+ = 0.04 indicates that the probability of the effect being positive is only 0.04, so that the probability of it being negative is 0.96.

### Brain responses

#### Main effect of valence: voxelwise analysis

Evidence for a main effect of valence was detected bilaterally in occipitotemporal regions linked to visual processing, right cingulate and peri-cingulate cortex, left insula, right parietal and bilaterally in frontal areas (Table 1 and Fig. 4.). Subcortically, valence effects were detected in the left cerebellum, left amygdala, and bilateral putamen.

**Table 1:** Main effect of valence. Thresholded at a voxel-level alpha value of 0.01 (FDR corrected).

| REGIONS | TALAIRACH<br>COORDINATES |  |  | CLUSTER<br>SIZE | F |
| --- | --- | --- | --- | --- | --- |
|  | X | Y | Z |  |  |
| Right Middle Temporal Gyrus (EBA) | 44 | -67 | -7 | 301 | 37.3 |
| Left Middle Temporal Gyrus (EBA) | -46 | -64 | -1 | 268 | 41.9 |
| Left Superior Frontal Gyrus ( dmPFC) | -4 | 50 | 32 | 228 | 32.8 |
| Left Putamen | -13 | 8 | -1 | 119 | 30.1 |
| Left Lingual Gyrus | -16 | -82 | -13 | 86 | 49.1 |
| Right Putamen/pallidum | 20 | 14 | -4 | 72 | 23.2 |
| Right Supplementary Motor Area | 11 | 8 | 56 | 62 | 36.3 |
| Right Middle Cingulate Cortex | 5 | 17 | 29 | 57 | 32.8 |
| Right Precentral Gyrus (BA 6/BA 4) | 38 | -7 | 38 | 57 | 28.9 |
| Right Inferior Frontal Gyrus (p. triang/vlPFC) | 50 | 38 | 8 | 41 | 24.3 |
| Right Superior Parietal Lobule/Precuneus | 26 | -49 | 44 | 24 | 22.9 |
| Left Insula | -34 | 26 | 8 | 15 | 18.9 |
| Left Precentral Gyrus (BA 4) | -43 | -7 | 38 | 15 | 20.1 |
| Right Precuneus (BA 7) | 11 | -58 | 47 | 13 | 20.2 |
| Left Cerebellum | -4 | -55 | -37 | 9 | 21.0 |
| Left Amygdala | -25 | -4 | -10 | 8 | 17.4 |

**Fig. 4.**
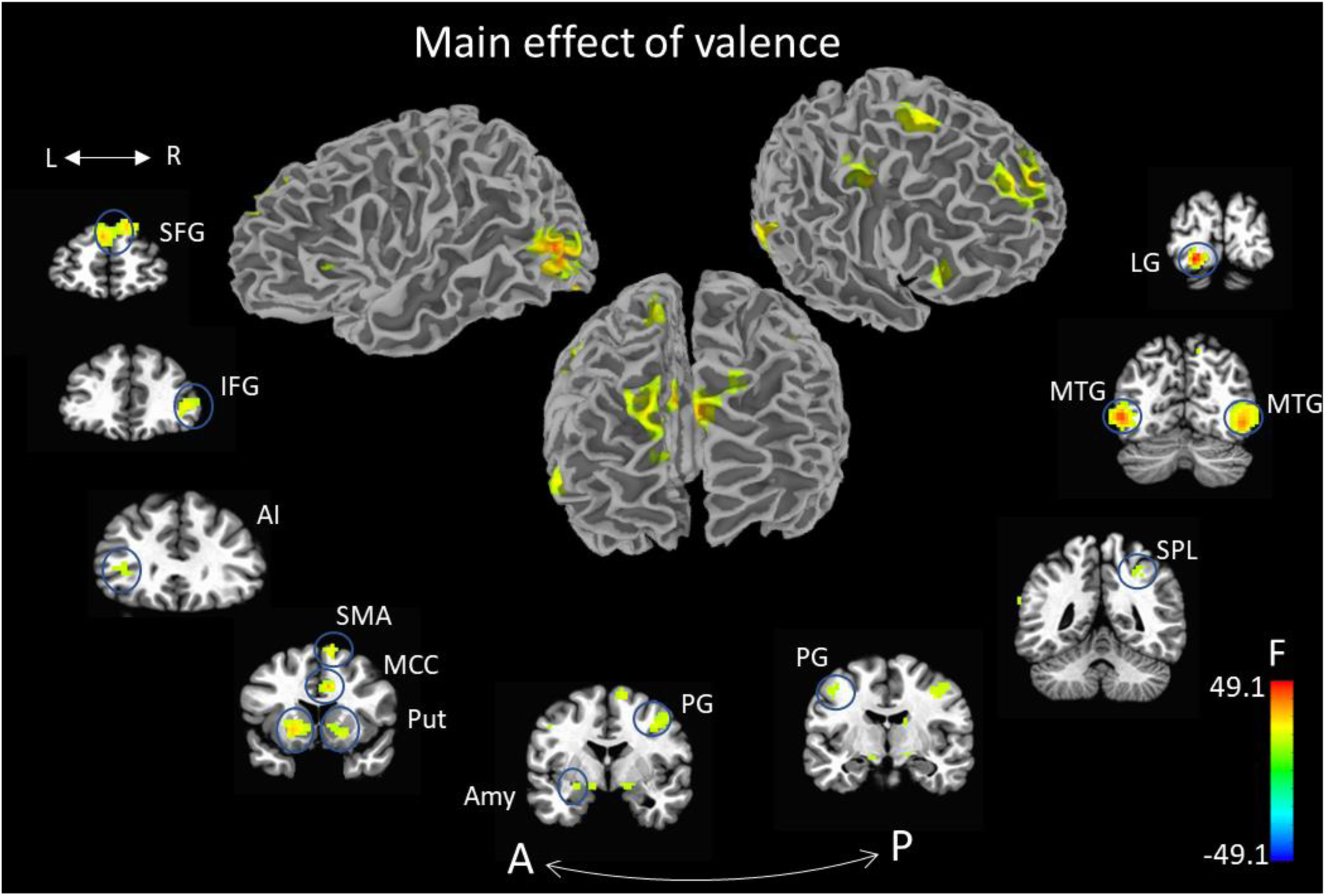
Voxelwise analysis: Main effect of valence. See Table 1 for the Tailarach coordinates. The color bar represents the value of the F-statistic. Abbreviations: A: Anterior; AI: Anterior insula; Amy: Amygdala; IFG: Inferior frontal gyrus; L: Left; LG: Lingual gyrus; MCC: Middle cingulated cortex; MTG: Middle temporal gyrus; P: Posterior; PG: Precentral gyrus; Put: Putamen; R: Right; SFG: Superior frontal gyrus; SMA: Supplementary motor area; SPL: Superior parietal lobule.

For completeness, the main effect of context is reported in Table 2.

**Table 2:** Main effect of context. Thresholded at a voxel-level alpha value of 0.01 (FDR corrected).

| REGIONS<br>MAIN EFFECT OF CONTEXT | TALAIRACH<br>COORDINATES |  |  | CLUSTE<br>R SIZE | F |
| --- | --- | --- | --- | --- | --- |
|  | X | Y | Z |  |  |
| Right Parahippocampal Gyrus | 26 | -43 | -7 | 3595 | 150.7 |
| Right Lingual Gyrus | 26 | -88 | -4 | 512 | 107.3 |
| Right Superior Parietal Lobule | 26 | -58 | 53 | 416 | 36.5 |
| Left Fusiform Gyrus | -37 | -76 | -13 | 369 | 74.9 |
| Right Middle Temporal Gyrus (EBA) | 41 | -67 | 17 | 276 | 79.4 |
| Left Superior Parietal Lobule | -28 | -52 | 47 | 100 | 20.9 |
| Left Precuneus | -1 | -55 | 38 | 74 | 23.4 |
| Left Parahippocampal Gyrus | -13 | -10 | -19 | 8 | 15.6 |

#### Valence by context interaction: voxelwise analysis

Although our goal was to investigate interaction effects in ROIs exhibiting evidence of a valence effect, for completeness we also performed a voxelwise analysis of the interaction, which was detected in visual areas (bilateral inferior/middle occipital gyrus, bilateral lingual gyrus, left inferior temporal/occipital gyrus, right middle occipital/temporal gyrus), as well as left inferior frontal gyrus (pars Opercularis), and right supplementary motor area (Table 3 and Fig. 5).

**Table 3:** Valence by context interaction in voxelwise analysis. Thresholded at a voxel-level alpha value of 0.05 (FDR corrected).

| REGIONS<br>VALENCE BY CONTEXT INTERACTION | TALAIRACH<br>COORDINATES |  |  | CLUSTE<br>R SIZE | F |
| --- | --- | --- | --- | --- | --- |
|  | X | Y | Z |  |  |
| Right Inferior/Middle Occipital Gyrus | 44 | -68 | -9 | 131 | 35.3 |
| Right Lingual Gyrus | 11 | -80 | -6 | 44 | 31.4 |
| Left Lingual Gyrus | -10 | -83 | -3 | 39 | 39.4 |
| Left Inferior/Middle Occipital Gyrus | -40 | -80 | -6 | 35 | 21.7 |
| Left inferior Frontal Gyrus (Opercularis) | -49 | 19 | 6 | 20 | 22.4 |
| Right Supplementary Motor Area | 2 | 10 | 54 | 15 | 18.7 |
| Left Inferior Temporal Gyrus | -40 | -50 | -15 | 13 | 21.3 |
| Right Superior Occipital Gyrus | 29 | -83 | 30 | 6 | 17.8 |

**Fig. 5.**
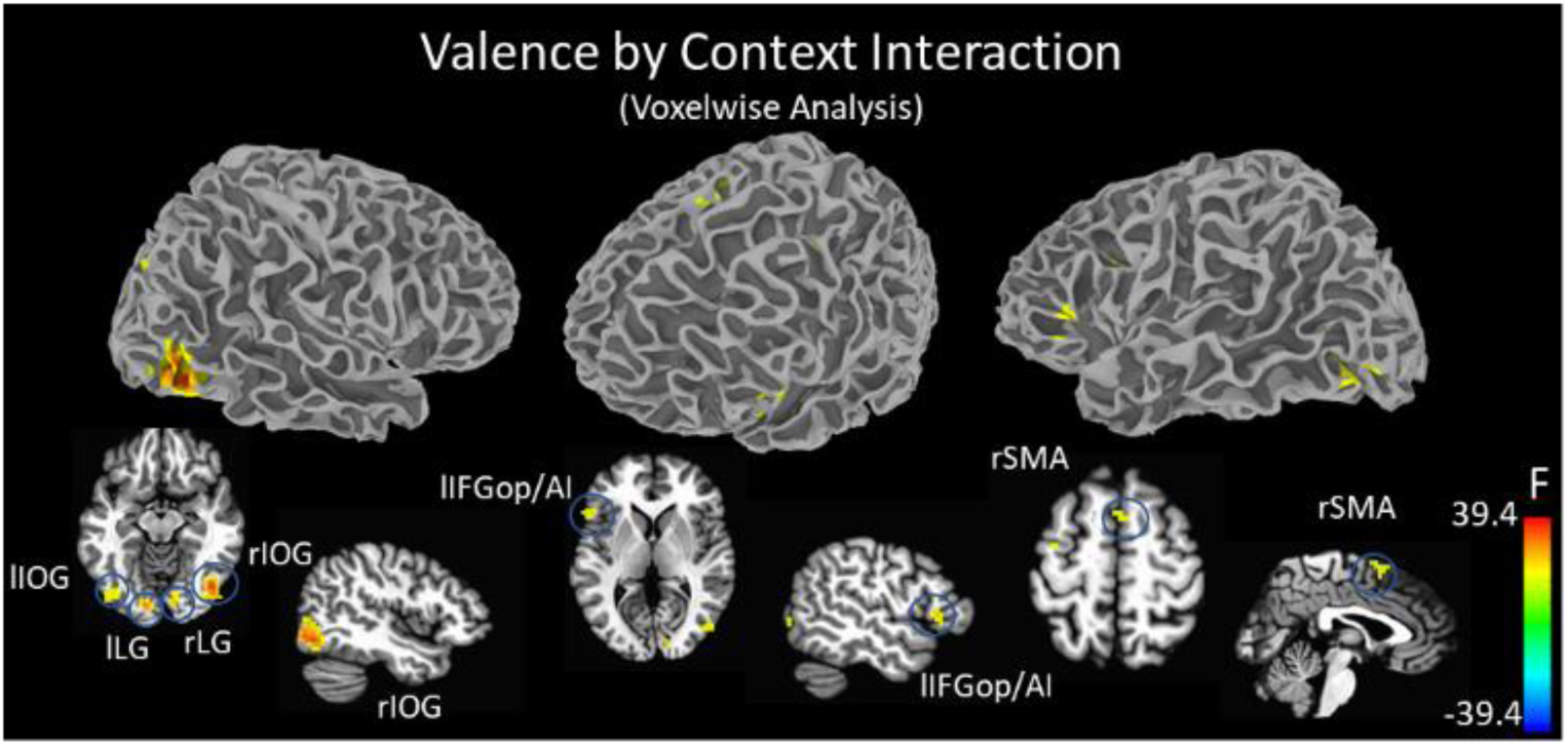
Voxelwise analysis: Valence by context interaction effect. See Table 3 for the Tailarach coordinates. The color bar represents the value of the F-statistic. Abbreviations: IFGop/AI: Inferior frontal gyrus pars opercularis / anterior insula; IOG: Inferior occipital gyrus; L: Left; LG: Lingual gyrus; R: Right; SFG: Superior frontal gyrus; SMA: Supplementary motor area.

#### Interaction between valence and context: Region of interest analysis

As stated, as our focus was to investigate whether or not threat direction influenced emotional modulation, interactions between valence and context were further investigated in ROIs selected based on the main effect of valence (Table 1). In all regions exhibiting evidence for a main effect of valence, activity was greater during threat conditions relative to neutral ones (except for the lingual gyrus, in which the reverse was observed).

We performed a Bayesian multilevel analysis that included all ROIs simultaneously in a single model (Methods). The posterior distributions for the valence by context effect are shown in Figure 6, where the color indicates P+, the probability of the effect being greater than zero. For example, strong evidence was determined for the middle temporal gyrus bilaterally (at sites labelled in the literature as the “extrastriate body area”), the right supplementary motor area (SMA), the left anterior insula, and the right precentral gyrus. But note that since the Bayesian framework is not dependent on a target false-positive rate (say, 0.05), we can consider all effect strengths in a continuous fashion. For completeness, Table 4 summarizes effect sizes (Cohen’s d) of the contrasts between threat and neutral valence for each ROI and context. Figure 7 shows the mean parameter estimate for each subject per condition.

**Table 4:**
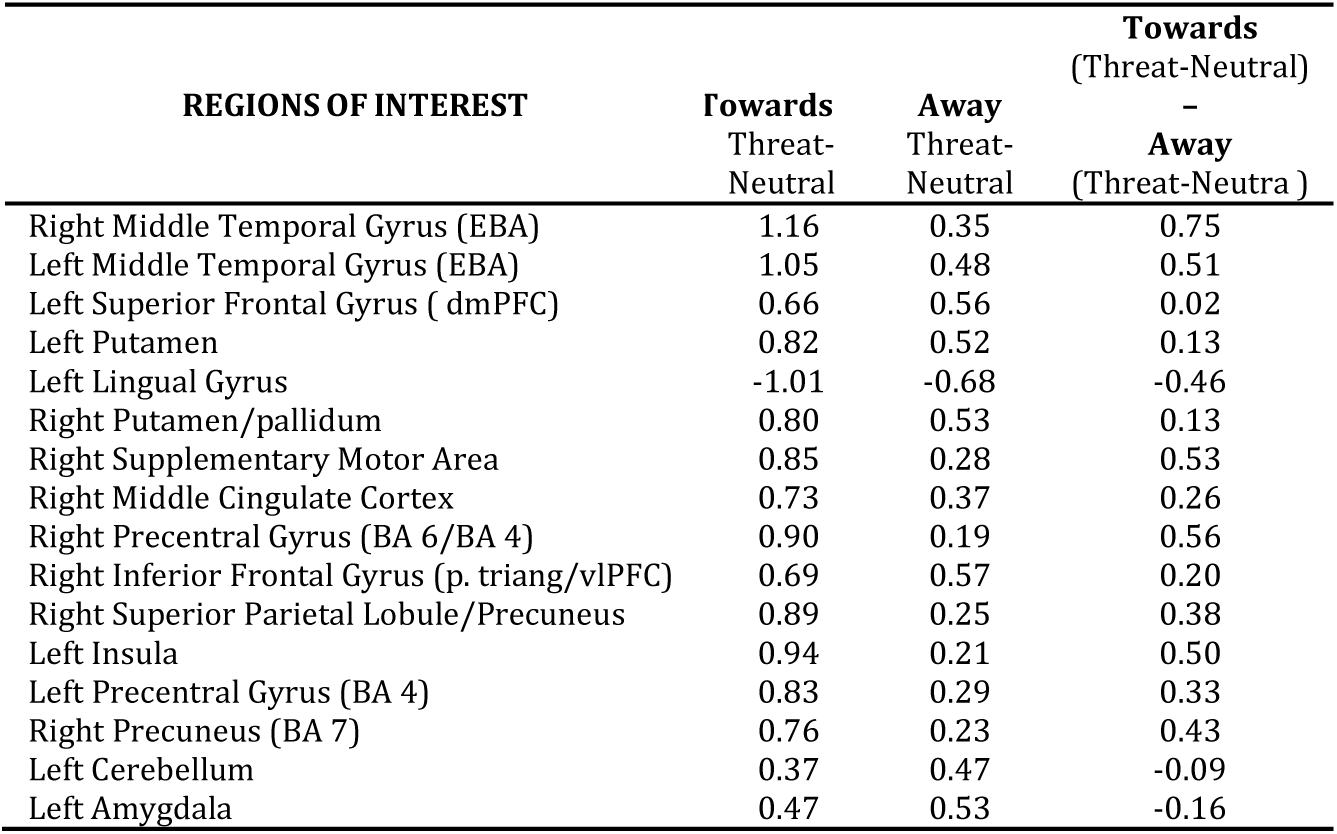
Effect size (Cohen’s D) of the difference between threat and neutral valence for each context and for the interaction. Talairach coordinates for each ROI are reported in Table 1.

**Fig. 6.**
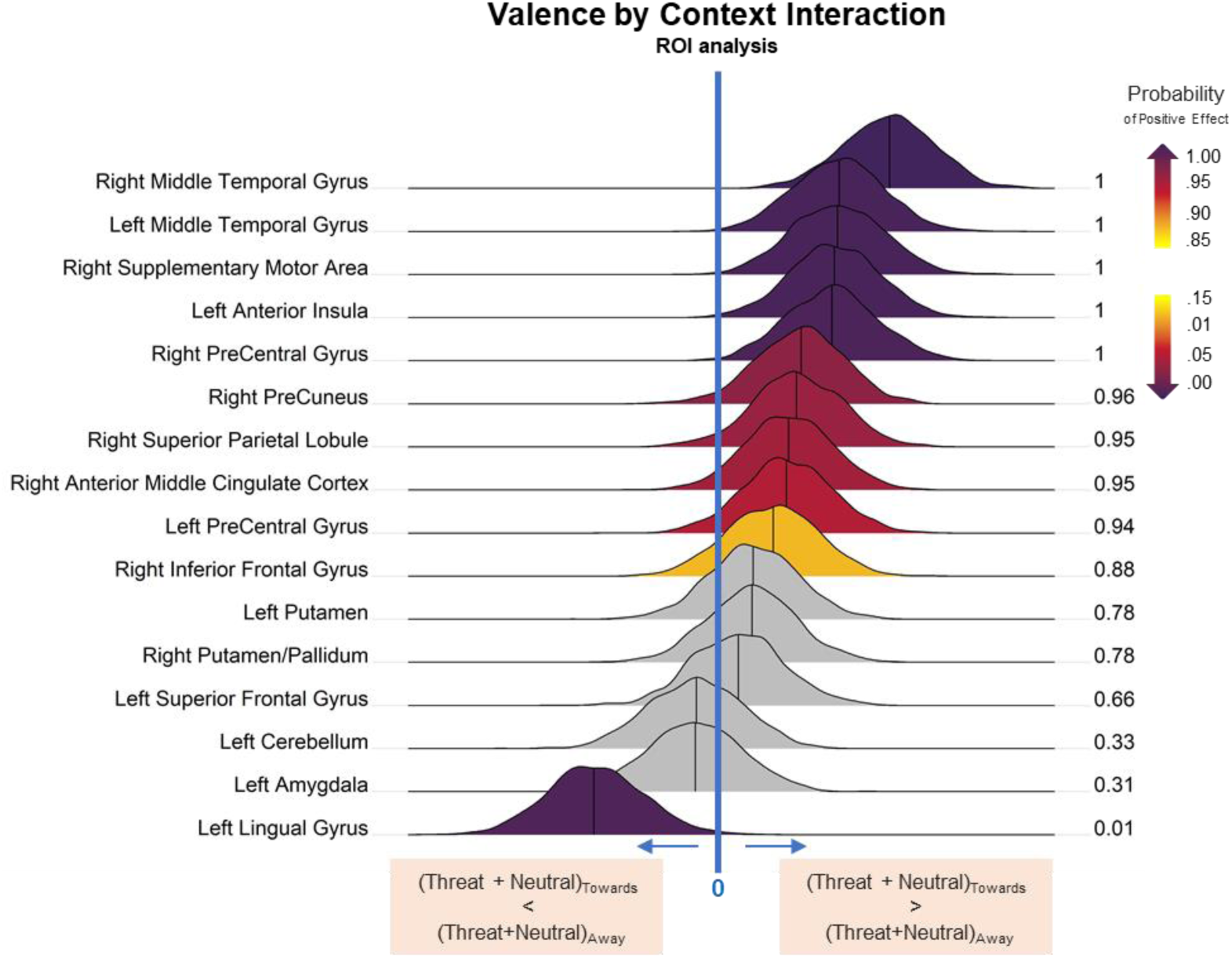
Bayesian posterior density plots of the valence by context interaction for ROIs reported in Table 1. The color bar indicates P+, the probability of the effect being greater than zero. Larger values of P+ convey support that the activity was greater for (Threat - Neutral)_Towards_ relative to (Threat - Neutral)_Away_; small values of P+ convey support of the reverse pattern. The color bar represents P+. See the caption of Figure 3 for discussion of interpretation.

**Fig. 7.**
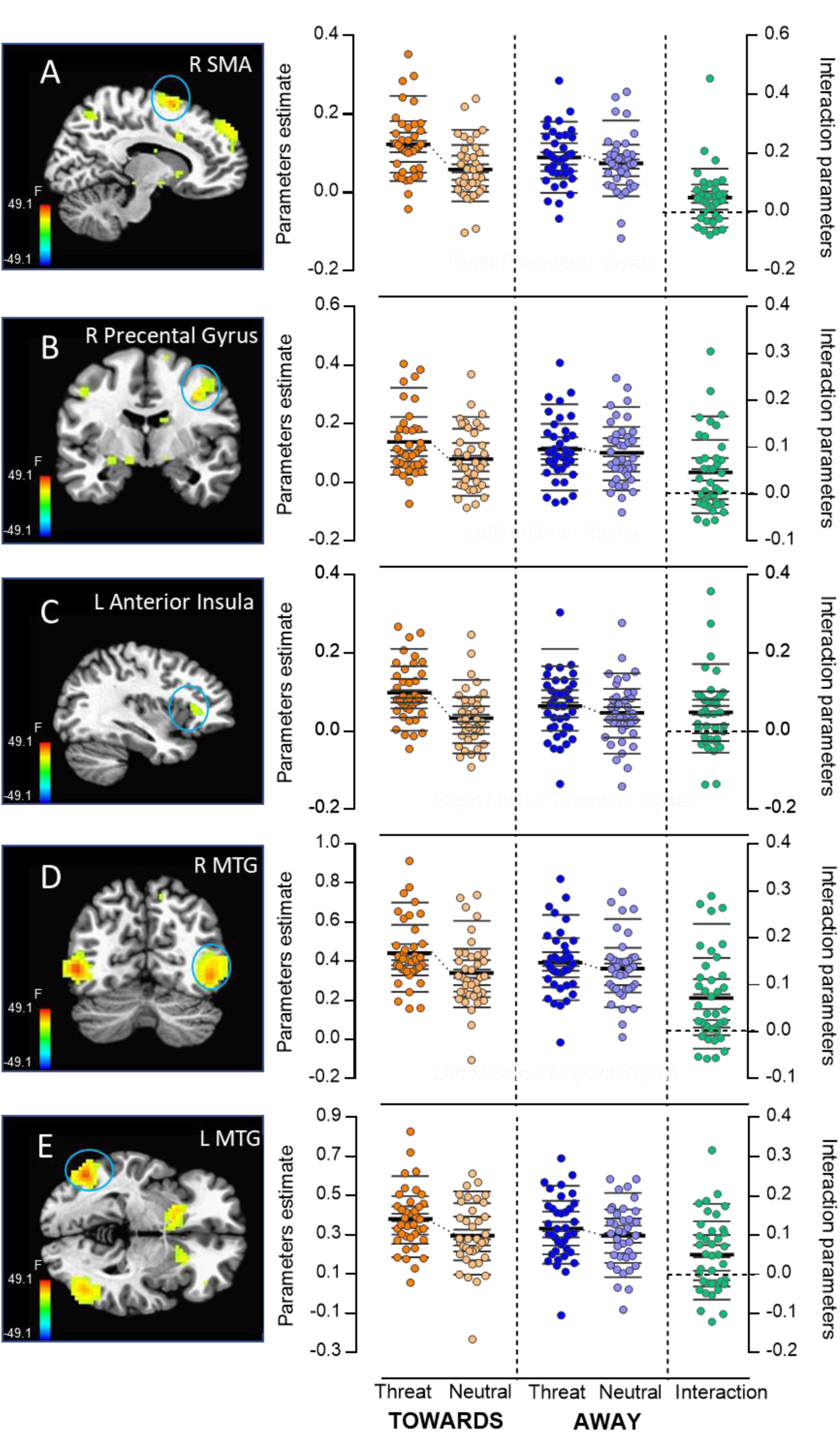
Regions of interest and valence by context interaction. (A) Supplementary Motor Area (11, 8, 56) (B) Precentral Gyrus (38, -7, 38) (C) Left Anterior Insula (−34, 26, 8) (D) Right Middle temporal Gyrus (MTG, Right: 44, -67, -7) (E) Left Middle temporal Gyrus (MTG, -46, -64, -1). Circles represent the mean parameter estimate for each subject per condition: Threat Towards, Neutral Towards, Threat Away, Neutral Away. The interaction index was calculated as follows: ((Threat - Neutral)_Towards_ - (Threat - Neutral)_Away_). Thicker lines mark the mean and thinner lines the deciles for each experimental condition. The thin purple line marks the shift between means. Data illustrations used some of the graphical tools proposed by Rousselet et al (Rousselet, Pernet & Wilcox, 2017).

#### Response pattern in the anterior MCC

We analyzed responses in terms of valence, context, and their interaction in the aMCC based on anatomical atlas-based ROIs. Bayesian analysis revealed strong evidence for the interaction on the left (P+ = 0.975), and good support on the right (P+ = 0.941). Both right and left were rather robustly driven by valence (left: P+ = 0.998; right: P+ = 0.998), but not by context (left: P+ = 0.449; right: P+ = 0.580). See Figure 8B for bayesian posterior density plots of interaction, valence and context effects and Figure 8C for mean parameter estimate for each subject per condition.

**Fig. 8.**
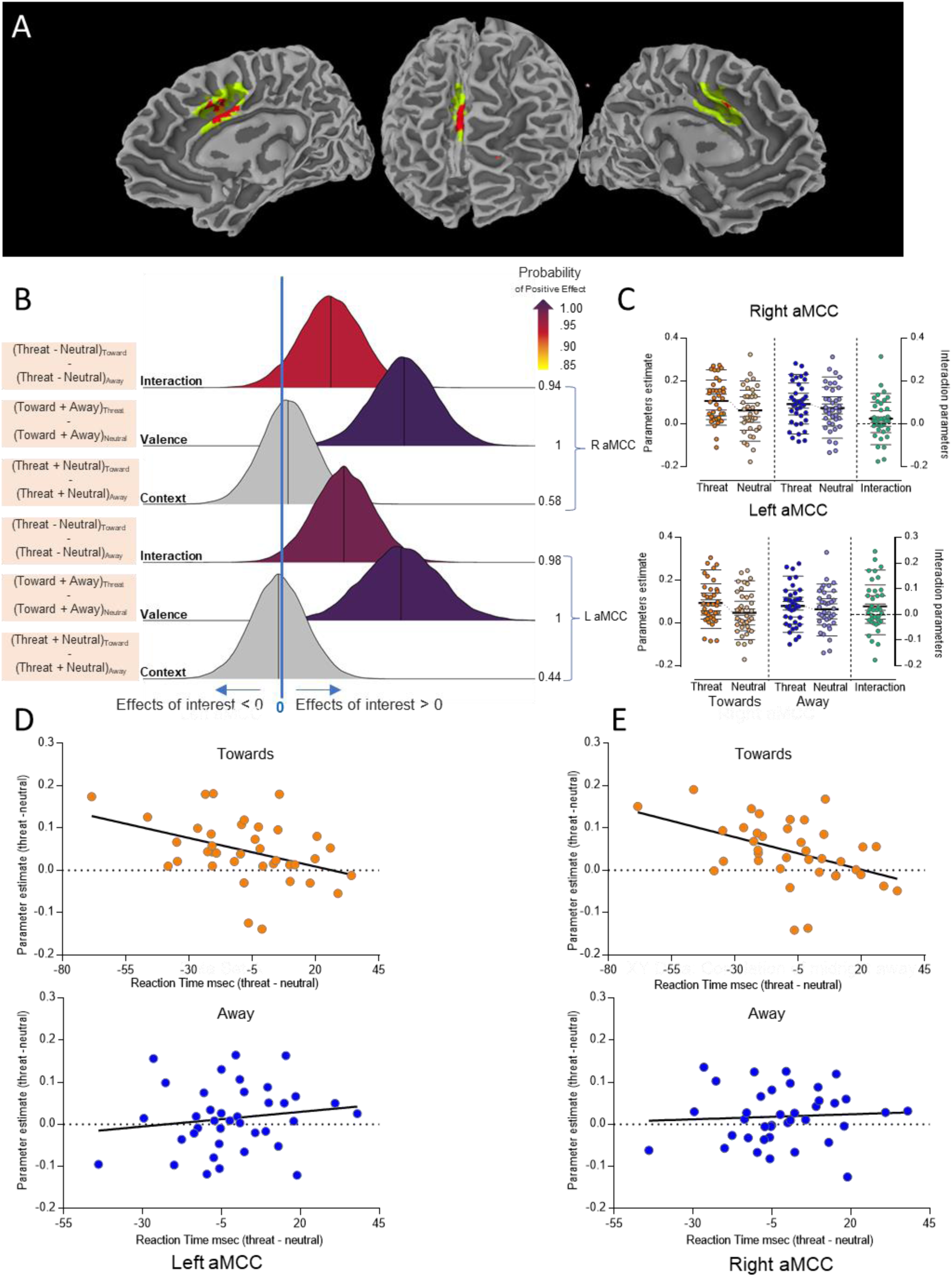
Anterior middle cingulate cortex (aMCC). (A) The anatomical ROI (green) is shown together with the midcingulate cluster that exhibited a main effect of valence (orange); the overlap between the two is shown in red. (B) Bayesian posterior density plots of interaction ((Threat - Neutral)_Towards_ - (Threat-Neutral)_Away_), valence ((Towards + Away)_Threat_ - (Towards + Away)_Neutral_) and context ((Threat + Neutral)_Towards_ - (Threat + Neutral)_Away_) effect for anterior right and left middle cingulate ROI based on atlas-based anatomical ROIs. The color bar indicates P+, the probability of the effect being greater than zero. The color bar represents P+. See the caption of Figure 3 for discussion of interpretation. (C) Parameter estimates for the anatomical aMCC ROI. (see Fig. 7 for additional definitions). (D-E) Brain-behavior correlations in the directed towards (top) and in the directed away context (bottom) for the left (D) and right (E) aMCC.

#### Brain-behavior correlations

A previous study by our group showed that aMCC activity paralleled behavioral emotional modulation in an aversive context (Pereira et al., 2010). Accordingly, we employed a behavioral modulation index defined as the mean RT to targets during threat trials minus the mean RT to targets during neutral trials, separately for the directed towards and directed away contexts. Corresponding indices were defined for aMCC activity, and a linear association between the two was evaluated. Figure 8 A illustrates the left and right anatomical aMCC ROI.

For the directed towards context, we observed evidence for a negative relationship between behaviour and brain responses (Figures 8 D and E, top): faster RTs for threat relative to neutral stimuli were associated with increased aMCC responses (threat vs. neutral). For the directed towards context, evidence was strong for the right (P+ = 0.029; Spearman rho = -0.47) and especially the left (P+ = 0.007; Spearman rho = -0.39) hemisphere. For the directed away context, evidence of a relationship between brain and behavior was not particularly noteworthy (left: P+ = 0.810; Spearman rho = 0.17; right: P+ = 0.643; Spearman rho = 0.11). A direct comparison of the association between the two contexts (towards vs. away) revealed strong support only for the right hemisphere (left: P+ = 0.212; right: P+ = 0.026).

We employed the same approach above to conduct exploratory analyses in regions showing some evidence of an interaction between valence and context (“some evidence” was defined as a P+ of approximately 0.95). Based on the posterior distributions in Fig. 6, we considered the following ROIs to investigate brain-behavior correlations: right and left middle temporal gyrus, right supplementary motor area, left anterior insula, right precentral gyrus, right precuneus, right superior parietal lobule and right aMCC. Posterior distributions for brain-behavior correlations for towards and away contexts and for the difference of the correlations between contexts are shown in Fig. 9. In the directed towards context, evidence of a brain behaviour correlation was especially strong in right aMCC but also evident in right SMA, left anterior insula, and right precentral gyrus. For the directed away context, brain-behaviour associations did not receive support from the data. For the explicit comparisons of correlations between towards and away contexts, there was weak support for the right aMCC, right SMA, and left insula. For completeness, Table 5 summarizes effect sizes (rho values) of the brain-behavior correlations for each ROI and context.

**Table 5:** Brain-behavior correlations for regions of interest with evidence of a valence by context interaction effect.

|  | TOWARDS<br>CONTEX | AWAY<br>CONTEXT |
| --- | --- | --- |
| REGIONS OF INTEREST | Rho | Rho |
| Right Middle Temporal Gyrus | 0.17 | 0.08 |
| Left Middle Temporal Gyrus | 0.06 | 0.24 |
| Right Supplementary Motor Area | -0.34 | 0.14 |
| Left Insula | -0.31 | 0.21 |
| Right Precentral Gyrus | -0.32 | 0.02 |
| Right Precuneus | -0.16 | 0.10 |
| Right Superior Parietal Lobule/Precuneus | -0.14 | -0.01 |
| Right Anterior Midcingulate cortex | -0.46 | 0.14 |

**Fig. 9.**
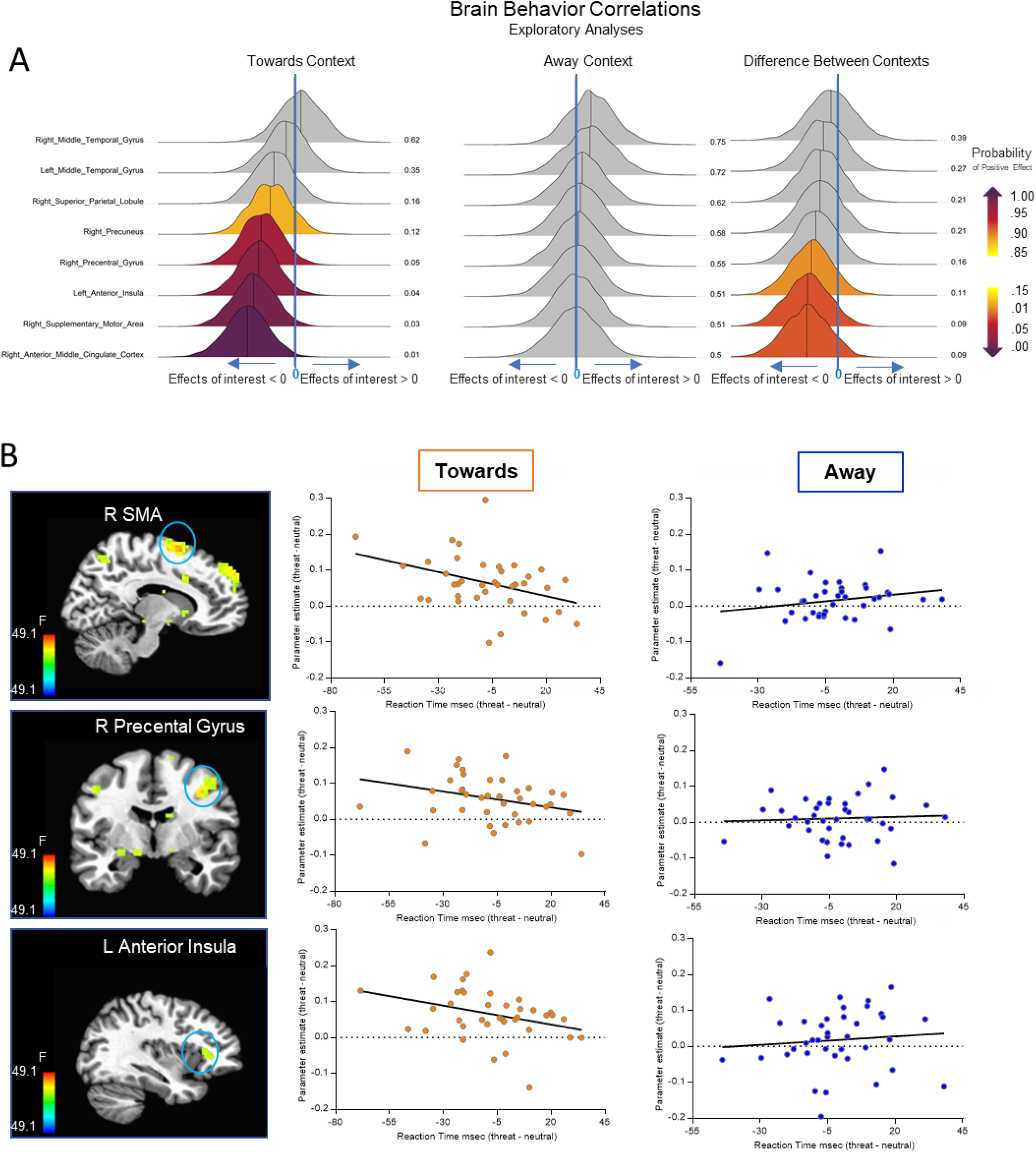
Brain-behavior relationship in regions showing evidence of an interaction between valence and context. (A) Bayesian posterior density plots of the brain-behavior correlation for the towards context, the away context, and the difference between them. The color bar indicates P+, the probability of the effect being greater than zero. See the caption of Figure 3 for discussion of interpretation. (B) Brain-behavior correlation in the directed towards (left plots) and in the directed away contexts (right plots) for the supplementary motor area (top), precentral gyrus (middle) and anterior insula (bottom).

## Discussion

In the present study, we sought to investigate how emotional processing is affected by context. In particular, we hypothesized that self-relevance would influence both behavior and associated brain responses. Self-relevance was manipulated by having participants view pictures containing guns that were directed at or away from them. We identified a group of brain areas where activity increased during threat relative to neutral conditions, with a subset of them (occipital cortex, precentral gyrus, supplementary motor area, and insula) exhibiting stronger evidence of a valence by context effect. Finally, we observed a brain-behavior correlation in the right and left aMCC, and to some extent in the right SMA, left anterior insula, and right precentral gyrus, indicating a closer relationship between evoked responses and target detection times during the self-relevant condition.

One of the features of the present work was that, wherever possible, it followed a Bayesian data analysis approach. Accordingly, we did not consider probability values dichotomously (say, “significant” if less than 0.05) but as providing continuous evidence for support for the hypothesis of interest, given the data. Furthermore, we adopted wide, poorly informative prior distributions that had very little discernible impact on our conclusions.

### Regions modulated by threat relevance

We observed robust valence by context interactions in regions involved in motor-related processing. Specifically, in both the right SMA and the right precentral gyrus, the differential response to threat vs. neutral was enhanced when the context was more relevant to the participant (directed towards condition). Whereas an increase in activation of cortical motor-related areas while participants observe negative stimuli has been reported (de Gelder et al., 2004; Grèzes, Pichon & de Gelder, 2007, Pichon, de Gelder & Grèzes, 2008, Pichon, de Gelder & Grèzes, 2009, Ahs et al., 2009, Pereira et al., 2010, Van den Stock et al., 2011, Conty et al., 2012, Pichon, de Gelder & Grèzes, 2012, Kveraga et al., 2015), the role of self-relevance remains poorly understood. Grèzes et al., (2013) reported that both the SMA and the precentral gyrus were engaged preferentially to body expressions of anger oriented to self when compared to anger oriented to other, and suggested that the recruitment of these areas might be related to the need of selecting specific behavioural strategies when one is the potential target of someone’s anger. Conty et al. (2012) combined fMRI and electroencephalography in humans, and obtained evidence that, 200 ms after stimulus onset, the premotor cortex integrated gaze, gesture, and emotion. They suggested that the early binding of visual social signals displayed by an agent engaged the dorsal pathway and the premotor cortex, possibly to facilitate the preparation of an adaptive response to another person’s immediate intention. We thus suggest that, in the context of our experiment, an increase in threat relevance impacts motor-related processing, possibly to implement an appropriate defensive response.

Another region that revealed a robust interaction effect between valence and context was in the middle temporal gyrus, at a location corresponding to the extrastriate body area, or EBA (Downing et al. 2001; Urgesi et al., 2004). Middle temporal gyrus activity was increased for threat stimuli in both contexts but the impact was greater when stimuli were directed towards the participant. The EBA site in the middle temporal gyrus has also been shown to be engaged by emotion-laden stimuli (Grosbras & Paus, 2006, Ponseti et al., 2006, Grèzes, Pichon & de Gelder, 2007; Peelen et al., 2007; Pichon, de Gelder & Grèzes, 2008; Flaisch et al., 2009; Sinke et al., 2010; Kret et al., 2011, Kveraga et al., 2015, Van den Stock et al., 2015). It has been proposed that the function of this area goes beyond the mere perception of body shape, as it might provide an interface between perceptual and motor processes (Astafiev et al., 2004; David et al., 2007; Kuhn et al., 2011; Tomasino, Weiss & Fink, 2012; Limanowski, Lutti, & Blankenburg, 2014; Orgs et al., 2016; Simos et al., 2017). For example, Zimmermann et al. (2017) suggested that the middle temporal gyrus/EBA interacts more strongly with dorsal-stream regions, when compared to other portions of the occipito-temporal cortex involved in processing body parts and object identification, and proposed that the area contributes to planning goal-directed actions.

### Anterior midcingulate cortex and behavioral modulation

We previously suggested that the aMCC is a site of interaction of negative valence and motor-related signals (Pereira et al., 2010), which motivated the examination here of self-relevance by using an anatomically based ROI. Our results revealed increased aMCC activity for threat vs. neutral stimuli in both directed towards and directed away threat contexts, consistent with an extensive literature pointing to the engagement of the aMCC in aversive processing (for a review see Vogt, 2005).

There has been growing interest in considering the midcingulate cortex as a unique cingulate region with a particular functional profile (Vogt el al 2016). Initial models of the anterior cingulate cortex suggested that rostral ACC and aMCC (also called dorsal ACC in some studies) specialize according to emotional and cognitive processes, respectively (Bush et al., 2000). In subsequent work, the aMCC has been proposed to be involved in emotion-, cognition-, and pain-related processing (Shackman et al., 2011); see also Misra & Coombes (2015). In addition, meta-analysis studies have proposed that the aMCC plays a central, integrative role in emotion regulation (Kohn et al, 2014), and is part of a core system for implementing self-control across emotion and action domains (Langner et al., 2018). In our previous study, the aMCC was recruited robustly only when participants performed a task in negative contexts, and responses mirrored the pattern of behavioral modulation associated with negative stimuli (Pereira et al., 2010). In the present study, we found a correlation between aMCC activity and behavior during the self-relevant threat context, whereby faster reaction times for threat (vs. neutral) stimuli were associated with increased activity in the aMCC (threat vs. safe). In other words, participants that exhibited greater speeding-up in RT for threat also exhibited increased threat-related brain responses. We also note that the reduction in RT for threat observed in the present study is in line with our previous observation that participants were quicker to perform a task in the presence of threat directed towards the self (Fernandes et al., 2013).

We conjecture that participants with greater recruitment of midcingulate cortex exhibited an enhanced implementation of active defensive responses. In line with this possibility, Straube (2009) found activation in the aMCC to be relevant when executive functions are more strongly necessary, such as in situations of high vigilance and action prompting. The present results also converge with the proposal that the aMCC is an important node for emotion and motor interaction in the service of adaptive responses to threat (Pereira et al., 2010, Shackman et al., 2011).

We note that a brain-behavior correlation analogous to that found for the aMCC was observed in the right SMA, precentral gyrus, and anterior insula. In this context, it is relevant that neurons in the aMCC (specifically in the rostral cingulate zone) project to other motor-related areas, including the premotor and supplementary motor cortices (Picard & Strick, 1996, see Shackman et al., 2011 for a review). These brain-behavior correlations indicate a closer relationship between evoked brain responses and target detection times during the self-relevant condition. In this context, and as noted above, Langner et al. (2018) proposed that the MCC, medial premotor regions, and anterior insula are essential for implementing self-control across emotion and action. In addition, a recent meta-analysis uncovered a potential association between the integrity of these regions and diverse psychiatric syndromes, including mood and anxiety disorders (Goodkind et al., 2015).

### Brain areas responding to threat

Of the areas exhibiting a main effect of valence, we would like to draw attention to the amygdala, a region that has been exhaustively studied in the context of negative processing. This region did not exhibit signs of a valence by context interaction; it did not seem to be affected by increasing threat relevance to the participant. This finding is relevant in the context of the debate of the nature of amygdala responses, namely the extent to which it is modulated by high-level factors such as attention and context, for example. Whereas we have reported that the amygdala is modulated by attention and contextual information (Pessoa, 2013), amygdala responses in the present study were not robustly sensitive to self-relevance. The present results illustrate how the amygdala functions, in some instances, in a more “basic” and less integrative manner.

### Limitations

Some limitations of the present work deserve discussion. First, we did not observe clear support for a valence by context interaction pattern in the RT data. Whereas we observed a reduction in RT when performing a task in the directed towards context in our previous behavioral study (Fernandes et al., 2013), evidence for this modulation effect was modest here. This difference is possibly due to the smaller sample size used in the current fMRI study, which also compromises the statistical power to detect such interaction effects in a voxelwise analysis. Second, scanning was performed at 1.5 Tesla, which has reduced sensitivity compared to 3 Tesla systems. In fact, we only detected a small amygdala cluster (8 voxels) with evidence of a main effect of emotion.

### Conclusions

How does the brain integrate emotion-related information and context? In the present study, we manipulated the self-relevance of threat stimuli by manipulating the direction of threat (guns pointed towards or away from the self). Using a Bayesian analysis framework, we identified multiple brain regions sensitive to threat direction, including the middle temporal gyrus/EBA, SMA, and precentral gyrus. We believe the context-sensitive recruitment of motor-related areas (SMA and precentral gyrus) is particularly noteworthy as they provide clues as to how emotion and action signals are integrated. In addition, the present results strengthen the idea that the aMCC is an important node for emotion and motor interaction, as responses in this brain region were correlated with behavior. More broadly, in the same manner that emotion interacts with perception and cognition, it is likely to interact with motor functions in the service of adaptive behaviors.

## Acknowledgments

This work was supported in part by federal and state Brazilian research agencies (CAPES and FAPERJ number 202.773/2016) to Liana Portugal. Luiz Pessoa received support from the National Institute of Mental Health (R01 MH071589).

## SUPPLEMENTAL MATERIAL

### Psychophysiological responses and evaluative characterization of stimuli (supplemental material)

Data of all stimuli characterization reported below was collected from independent samples.

### Valence and arousal ratings

Following the protocol developed by Lang and colleagues (1997), the pictures were rated on a scale of 1–9 in terms of pleasure and arousal by a separate group of 134 participants (104 females, 21.5 years ± 3.36) using the paper-and-pencil version of the Self-Assessment Manikin (Bradley and Lang, 1994). The mean values of valence and arousal for each picture category are shown in Table S1.

### Complexity ratings

Another important aspect that we sought to match between the threat and neutral stimuli was picture complexity. This aspect was considered because a previous study showed that picture complexity (clear figure-ground pictures compared with complex scenes that depicted multiple objects) rather than emotionality was responsible for some of the differences observed in the recorded neural responses to neutral and emotional pictures (Bradley et al., 2007). We attempted to minimize this confounding factor by selecting only emotional and neutral stimuli with approximately the same level of complexity, i.e., they were all clear figure-ground pictures with the same perceptual properties. To assess the adequacy of this a priori selection, we followed the procedures of Bradley et al. (2007) and asked an independent sample of 58 students (42 female) to rate picture complexity on a scale of 1–9 (1=clear figure-ground, 9=complex scenes). The results corroborated our a priori selection of the pictures (Table S2).

### Skin conductance and threat perception ratings

We measured skin conductance response to evaluate the differential emotional impact of the pictures and collected evaluative ratings of the stimuli to check if increasing self-relevance of the stimuli would be associated to increased emotional impact.

Sixty one participants (42 females, 20.8 years, ± 2.8) were exposed to a subset (n=64, 16 from each from each category) of the pictures used in the fmri experiment. The experimental session consisted of two blocks: the order of the blocks was randomized, and the presentation of neutral and threat pictures within the experiment was pseudo-randomized among participants. Each trial began with a fixation cross that was presented for 6–8 s, which was followed by a picture presented for 6 s. The participants were instructed to observe the picture during its presentation. The skin conductance response (SCR) were collected with a GSR150 module coupled to the MP150 low-pass filtered (1 Hz), sampled at 500 Hz via BIOPAC (BIOPAC Systems Inc.). The analyses were implemented using the EDA toolbox (https://github.com/mateusjoffily/EDA/wiki, Matheus Joffilly). Therefore, skin conductance responses with amplitudes higher than 0.02 ms were analysed within a temporal window of 1 to 3 s after each picture onset. Finally, the number of skin conductance responses to the entire set of pictures was calculated for each condition (Oliveira et al., 2009). We performed repeated measures ANOVA with direction (block stimuli directed towards the observer and no direct block stimuli) and valence (emotional and neutral) as the within-subjects factors. ANOVAs demonstrated a significant main effect of valence (F(1,60)=17.191, p=0.0001), which revealed a greater number of skin conductance responses to threat compared with neutral stimuli (threat=1.56, neutral=1.03, p=0.002). We also observed a significant interaction between direction and valence (F(1, 60)=4.4419, p=0.039). Post hoc (Newman-Keuls) analyses revealed that threat stimuli directed towards the observer induced a greater number of skin conductance responses compared to all conditions (1.82, SD=1.68). This result suggests that this manipulation was successful: threat stimuli directed towards the observer promoted greater activation of autonomic response, corroborating the idea of prompting of differential defensive reactions.

After the skin conductance recording session, the same participants realized an evaluative session which consisted of the presentation of the same four sets of pictures followed by the respective ratings on the evaluative questionnaire. The evaluative questionnaire was answered using a paper and pencil on a Likert-like scale of 9 points. Questions were related to the dimensions of the threat perception: (i) threat magnitude, (ii) the distance between the threat and the participant (proximity), (iii) the escapability of the situation (inescapability) and (iv) the presence of available hiding places (impossibility of hiding).

Ratings obtained for all dimensions of the threat perception scale were summed, creating a threat perception index for threat stimuli directed towards and directed away from the observer. Then, the threat perception index for each block was compared using a two-tailed paired *t*-test, which revealed that the threat index for the directed towards block (18.56) was significantly greater (*t*(61) = 7.61, p < 0.0001) than the threat index for the directed away block (12.57). These data indicate that threat in the directed towards block was considered more intense, near and inescapable and that there was a reduced possibility of hiding. These results are in agreement to what was described in previous studies that used similar threat stimuli (Fernandes et al (2013?) and Bastos et al (2016?)).

---

**Table S1:** The mean values and standard deviations (SDs) for pleasure and arousal for each experimental condition.

|  | Threat towards |  | Neutral towards |  | Threat away |  | Neutral away |  |
| --- | --- | --- | --- | --- | --- | --- | --- | --- |
|  | Mean | SD | Mean | SD | Mean | SD | Mean | SD |
| Pleasure | 2.06 | 0.37 | 5.20 | 0,39 | 2.73 | 0.71 | 5.47 | 0.83 |
| Arousal | 6.56 | 0.70 | 3.65 | 0,56 | 5.50 | 1.20 | 3.63 | 1.32 |

**Table S2:** The mean values and standard deviations (SDs) for complexity ratings and physical features for each experimental condition.

|  | Threat towards |  | Neutral towards |  | Threat away |  | Neutral away |  |
| --- | --- | --- | --- | --- | --- | --- | --- | --- |
|  | mean | SD | Mean | SD | Mean | SD | Mean | SD |
| Complexity | 3.00 | 0.74 | 2.92 | 0.57 | 2.59 | 0,77 | 3,02 | 1,04 |
| Brightness | 76.47 | 23.75 | 79.33 | 24.98 | 92.84 | 36,86 | 87,07 | 37,10 |
| Contrast | 25.43 | 9.73 | 27.67 | 8.78 | 20.95 | 10,57 | 21,42 | 7,37 |

## References

Ahs, F., Pissiota, A., Michelgård, A., Frans, O., Furmark, T., Appel, L., & Fredrikson, M. (2009). Disentangling the web of fear: amygdala reactivity and functional connectivity in spider and snake phobia. Psychiatry Research, 172, 103–108. http://dx.doi.org/10.1016/j.pscychresns.2008.11.004

Astafiev, S. V., Stanley, C. M., Shulman, G. L., & Corbetta, M. (2004). Extrastriate body area in human occipital cortex responds to the performance of motor actions. Nature Neuroscience, 7, 542–548. http://dx.doi.org/10.1038/nn1241

Azevedo, T. M., Volchan, E., Imbiriba, L. A., Rodrigues, E. C., Oliveira, J. M., Oliveira, L. F., … Vargas, C. D. (2005). A freezing-like posture to pictures of mutilation. Psychophysiology, 42, 255–260. https://doi.org/10.1111/j.1469-8986.2005.00287.x

Bancaud, J., & Talairach, J. (1992). Clinical semiology of frontal lobe seizures. Advances in Neurology, 57, 3–58.

Bastos, A. F., Vieira, A. S., Oliveira, J. M., Oliveira, L., Pereira, MG, Figueira I., …Volchan, E. (2016). Stop or move: Defensive strategies in humans. Behavioural Brain Research, 302, 252–262. https://doi.org/10.1016/j.bbr.2016.01.043

Blanchard, R.J. & Blanchard, D.C. (1971). Defensive reactions in the albino rat, Learn.Motiv. 2, 351–362, http://dx.doi.org/10.1016/0023-9690(71)90016-6.

Blakemore, R.L. & Vuilleumier, P (2017). An Emotional Call to Action: Integrating Affective Neuroscience in Models of Motor Control. Emotion Review, 9 (4), 299–309 https://doi.org/10.1177/1754073916670020

Benjamin, D. J., Berger, J. O., Johannesson, M., Nosek, B. A., Wagenmakers, E. J., Berk, R., … Johnson, V. E. (2018). Redefine statistical significance. Nature Human Behaviour, 2 (1), 6–10. http://dx.doi.org/10.1038/s41562-017-0189-z.

Blanchard, R. J., Flannelly, K. J., & Blanchard, D. C. (1986). Defensive behaviors of laboratory and wild Rattus norvegicus. Journal of Comparative Psychology, 100, 101–107. http://dx.doi.org/10.1037/0735-7036.100.2.101

Bürkner, P.-C. (2017). brms: An R package for bayesian multilevel models using Stan. Journal of Statistical Software, 80 (1), 1–28. https://doi.org/10.18637/jss.v080.i01

Bush, G., Luu, P., & Posner, M. I. (2000). Cognitive and emotional influences in anterior cingulate cortex. Trends in Cognitive Sciences, 4, 215–222. https://doi.org/10.1016/S1364-6613(00)01483-2

Campagnoli, R. R., Krutman, L., Vargas, C. D., Lobo, I., Oliveira, J. M., Oliveira, L., … Volchan, E. (2015). Preparing to caress: a neural signature of social bonding. Frontiers in psychology, 6, 16. http://dx.doi.org/10.3389/fpsyg.2015.00016

Chen, G., Taylor, P. A., Cox, R. W., & Pessoa, L. (2019a). How to Deal with Multiplicity in Neuroimaging? A Case for Global Calibration. bioRxiv, 706747. https://doi.org/10.1101/706747

Chen, G., Xiao, Y., Taylor, P. A., Rajendra, J. K., Riggins, T., Geng, F., … & Cox, R. W. (2019b). Handling Multiplicity in Neuroimaging through Bayesian Lenses with Multilevel Modeling. Neuroinformatics, 1–31. http://dx.doi.org/10.1007/s12021-018-9409-6

Chen, M., & Bargh, J. A. (1999). Immediate behavioral predispositions to approach or avoid the stimulus. Pers. Soc. Psychol. Bull. 25, 215–224. http://dx.doi.org/10.1177/0146167299025002007

Coelho, C. M., Lipp, O. V., Marinovic, W., Wallis, G., & Riek, S. (2010). Increased corticospinal excitability induced by unpleasant visual stimuli. Neuroscience Letters, 481, 135–138. http://dx.doi.org/10.1016/j.neulet.2010.03.027

Cohen, M.S. (1997). Parametric analysis of fMRI data using linear systems methods. Neuroimage, 6, 93–103. http://dx.doi.org/10.1006/nimg.1997.0278

Conty, L., Dezecache, G., Hugueville, L., & Grèzes, J. (2012). Early binding of gaze, gesture, and emotion: neural time course and correlates. The Journal of Neuroscience, 32, 4531–4539. http://dx.doi.org/10.1523/JNEUROSCI.5636-11.2012.

Cox, R. (1996). AFNI: Software for analysis and visualization of functional magnetic resonance neuroimages. Computers and Biomedical Research, 29, 162–173. http://dx.doi.org/10.1006/cbmr.1996.0014

Damasio, A. R. (1999). The feeling of what happens: Body and emotion in the making of consciousness.

Darwin, C. (1872). The expression of the emotions in man and animals. London: J. Murray.

David, N., Cohen, M. X., Newen, A., Bewernick, B. H., Shah, N. J., Fink, G. R., & Vogeley, K. (2007). The extrastriate cortex distinguishes between the consequences of one’s own and others’ behavior. NeuroImage, 36, 1004–1014. http://dx.doi.org/10.1016/j.neuroimage.2007.03.030

de Gelder, B., Snyder, J., Greve D., Gerard, G., & Hadjikhani, N. (2004). Fear fosters flight: A mechanism for fear contagion when perceiving emotion expressed by a whole body. Proceedings of the National Academy of Sciences, 101, 16701-16706, 2004. http://dx.doi.org/10.1073/pnas.0407042101

de Oliveira, L. A., Imbiriba, L. A., Russo, M. M., Nogueira-Campos, A. A., Rodrigues Ede, C., Pereira, M. G., … Vargas, C. D. (2012). Preparing to grasp emotionally laden stimuli. PLoS One, 7, e45235. http://dx.doi.org/10.1371/journal.pone.0045235

Destrieux, C., Fischl, B., Dale, A., & Halgren, E. (2010). Automatic parcellation of human cortical gyri and sulci using standard anatomical nomenclature. NeuroImage, 53(1), 1–15. http://dx.doi.org/10.1016/j.neuroimage.2010.06.010

Downing, P. E., Jiang, Y., Shuman, M., & Kanwisher, N. G. (2001). A cortical area selective for visual processing of the human body. Science, 293, 2470–2473, 2001. http://dx.doi.org/10.1126/science.1063414

Facchinetti, L. D., Imbiriba, L. A., Azevedo, T. M., Vargas, C. D., & Volchan, E. (2006). Postural modulation induced by pictures depicting prosocial or dangerous contexts. Neuroscience Letters, 410 (1), 52–56. http://dx.doi.org/10.1016/j.neulet.2006.09.063

Fanselow, M. S., & Lester, L. S. (1988). A functional behavioristic approach to aversively motivated behavior: Predatory imminence as a determinant of the topography of defensive behavior. In R. C. Bolles & M. D. Beecher (Eds.), Evolution and learning (pp. 185–212). Hillsdale, NJ, US: Lawrence Erlbaum Associates, Inc.

Fernandes, O., Jr, Portugal, L. C., Alves, R. C., Campagnoli, R. R., Mocaiber, I., David, I. P., … Pereira, M. G. (2013). How you perceive threat determines your behavior. Frontiers in human neuroscience, 7, 632. http://dx.doi.org/10.3389/fnhum.2013.00632

Fernandes, O. Jr., Portugal, L. C. L., Alves, R. C. S., Arruda-Sanchez, T., Rao, A., Volchan, E., … Mourao-Miranda, J. (2017). Decoding negative affect personality trait from patterns of brain activation to threat stimuli. NeuroImage, 145, 337–345. http://dx.doi.org/10.1016/j.neuroimage.2015.12.050

Flaisch, T., Schupp, H. T., Renner, B., & Junghofer, M. (2009). Neural systems of visual attention responding to emotional gestures. NeuroImage, 45, 1339–1346. http://dx.doi.org/10.1016/j.neuroimage.2008.12.073

Frijda, N. H. (1986). Studies in emotion and social interaction. The emotions. New York, NY, US: Cambridge University Press; Paris, France: Editions de la Maison des Sciences de l’Homme.

Friston, K. J., Frith, C. D., Turner, R., & Frackowiak, R. S. J. (1995). Characterizing evoked hemodynamics with fMRI. NeuroImage, 2, 157–165. https://doi.org/10.1006/nimg.1995.1019

Grèzes, J., Pichon, S., & de Gelder, B. (2007). Perceiving fear in dynamic body expressions. NeuroImage, 35, 959–967. http://dx.doi.org/10.1016/j.neuroimage.2006.11.030

Grèzes, J., Philip, L., Chadwick, M., Dezecache, G., Soussignan, R., & Conty, L. (2013). Self-relevance appraisal influences facial reactions to emotional body expressions. PloS One, 8, e55885. http://dx.doi.org/10.1371/journal.pone.0055885

Goodkind M, Eickhoff SB, Oathes DJ, Jiang Y, Chang A, Jones-Hagata LB, Ortega BN, Zaiko YV, Roach EL, Korgaonkar MS, Grieve SM, Galatzer-Levy I, Fox PT, Etkin A. Identification of a common neurobiological substrate for mental illness. JAMA Psychiatry. 2015;72:305–315. http://dx.doi.org/10.1001/jamapsychiatry.2014.2206.

Grosbras, M.H., & Paus, T. (2006). Brain networks involved in viewing angry hands or faces. Cerebral Cortex, 16, 1087–1096. http://dx.doi.org/10.1093/cercor/bhj050

Hajcak, G., Molnar, C., George, M. S., Bolger, K., Koola, J., & Nahas, Z. (2007). Emotion facilitates action: a transcranial magnetic stimulation study of motor cortex excitability during picture viewing. Psychophysiology, 44 (1), 91–97. http://dx.doi.org/10.1111/j.1469-8986.2006.00487.x

Kohn, N., Eickhoff, S. B., Scheller, M., Laird, A. R., Fox, P. T., & Habel, U. (2014). Neural network of cognitive emotion regulation--an ALE meta-analysis and MACM analysis. NeuroImage, 87, 345–355. https://doi.org/10.1016/j.neuroimage.2013.11.001

Kolesar, T. A., Kornelsen, J., & Smith, S. D. (2017). Separating neural activity associated with emotion and implied motion: An fMRI study. Emotion, 17(1), 131–140. http://dx.doi.org/10.1037/emo0000209

Kornelsen, J., Smith, S. D., & McIver, T. A. (2014). A neural correlate of visceral emotional responses: evidence from fMRI of the thoracic spinal cord. Social cognitive and affective neuroscience, 10, 584–588. http://dx.doi.org/10.1093/scan/nsu092

Kret, M. E., Pichon, S., Grezes, J., & de Gelder, B. (2011). Similarities and differences in perceiving threat from dynamic faces and bodies. An fMRI study. NeuroImage, 54, 1755–1762. http://dx.doi.org/10.1016/j.neuroimage.2010.08.012

Krieglmeyer, R., & Deutsch, R. (2010). Comparing measures of approach-avoidance behaviour: The manikin task vs. two versions of the joystick task. Cognition and Emotion, 24, 810–828. https://doi.org/10.1080/02699930903047298

Kühn, S., Keizer, A., Rombouts, S. A., & Hommel, B. (2011). The functional and neural mechanism of action preparation: roles of EBA and FFA in voluntary action control. Journal of Cognitive Neuroscience, 23(1): 214–20. http://dx.doi.org/10.1162/jocn.2010.21418

Kveraga, K., Boshyan, J., Adams, R. B., Mote, J., Betz, N., Ward, N., … Barrett, L. F. (2015). If it bleeds, it leads: separating threat from mere negativity. Social Cognitive and Affective Neuroscience, 10 (1), 28–35. http://dx.doi.org/10.1093/scan/nsu007

Lang, P. J., Bradley, M. M., & Cuthbert, B. N. (1997). Motivated attention: Affect, activation, and action. In P. J. Lang, R. F. Simons, & M. T. Balaban (Eds.), Attention and orienting: Sensory and motivational processes (pp. 97–135). Mahwah, NJ, US: Lawrence Erlbaum Associates Publishers.

Lang, P. J., Bradley, M. M., & Cuthbert, B. N. (2005). International affective picture system (IAPS): affective ratings of pictures and instruction manual. Gainesville, FL: University of Florida.

Langner, R., Leiberg, S., Hoffstaedter, F., & Eickhoff, S. B. (2018). Towards a human self-regulation system: Common and distinct neural signatures of emotional and behavioural control. Neuroscience & Biobehavioral Reviews, 90, 400–410. https://doi.org/10.1016/j.neubiorev.2018.04.022

Limanowski, J., Lutti, A., & Blankenburg, F. (2014). The extrastriate body area is involved in illusory limb ownership. NeuroImage, 86, 514–524. http://dx.doi.org/10.1016/j.neuroimage.2013.10.035

McIver, T. A., Kornelsen, J., & Smith, S. D. (2013). Limb-specific emotional modulation of cervical spinal cord neurons. Cognitive, Affective & Behavioral Neuroscience, 13, 464–472. http://dx.doi.org/10.3758/s13415-013-0154-x

McShane, B.B., Gal, D., Gelman, A., Robert, C., & Tackett, J L. (2017). Abandon Statistical Significance. The American Statistician, 73, 235–245. https://doi.org/10.1080/00031305.2018.1527253

Meyer, C., Padmala, S., & Pessoa, L. (2019). Dynamic threat processing. Journal of Cognitive Neuroscience, 31, 522–542. https://doi.org/10.1162/jocn_a_01363

Misra, G., & Coombes, S.A. (2015). Neuroimaging Evidence of Motor Control and Pain Processing in the Human Midcingulate Cortex. Cerebral Cortex, 25, 1906–1919. https://doi.org/10.1093/cercor/bhu001.

Nogueira-Campos, A. A., de Oliveira, L. A., Della-Maggiore, V., Esteves, P. O., Rodrigues Ede, C., & Vargas, C. D. (2014). Corticospinal excitability preceding the grasping of emotion-laden stimuli. PLoS One, 9, e94824 http://dx.doi.org/10.1371/journal.pone.0094824

Oliveri, M., Babiloni, C., Filippi, M. M., Caltagirone, C., Babiloni, F., Cicinelli, P., … Rossini, P. M. (2003). Influence of the supplementary motor area on primary motor cortex excitability during movements triggered by neutral or emotionally unpleasant visual cues. Experimental Brain Research, 149, 214–221. http://dx.doi.org/10.1007/s00221-002-1346-8

Orgs, G., Dovern, A., Hagura, N., Haggard, P., Fink, G. R., & Weiss, P. H. (2015). Constructing Visual Perception of Body Movement with the Motor Cortex. Cerebral cortex (New York, N.Y.: 1991), 26(1), 440–449. http://dx.doi.org/10.1093/cercor/bhv262

Peelen, M. V., Atkinson, A. P., Andersson, F., & Vuilleumier, P. (2007). Emotional modulation of body-selective visual areas. Social cognitive and affective neuroscience, 2, 274–283. http://dx.doi.org/10.1093/scan/nsm023

Pereira, M. G., de Oliveira, L., Erthal, F. S., Joffily, M., Mocaiber, I. F., Volchan, E., & Pessoa, L. (2010). Emotion affects action: Midcingulate cortex as a pivotal node of interaction between negative emotion and motor signals. Cognitive, affective & behavioral neuroscience, 10(1), 94–106. http://dx.doi.org/10.3758/CABN.10.1.94

Pernet, C. R., Wilcox, R., & Rousselet, G. A. (2013). Robust correlation analyses: false positive and power validation using a new open source matlab toolbox. Frontiers in psychology, 3, 606. http://dx.doi.org/10.3389/fpsyg.2012.00606

Pessoa, L. (2013). The cognitive-emotional brain: From interactions to integration. Cambridge, MA, US: MIT Press.

Phaf, R. H., Mohr, S. E., Rotteveel, M., & Wicherts, J. M. (2014). Approach, avoidance, and affect: a meta-analysis of approach-avoidance tendencies in manual reaction time tasks. Frontiers in psychology, 5, 378. http://dx.doi.org/10.3389/fpsyg.2014.00378

Picard, N., & Strick, P. L. (1996). Motor areas of the medial wall: A review of their location and functional activation. Cerebral Cortex, 6, 342–353. http://dx.doi.org/10.1093/cercor/6.3.342

Pichon, S., de Gelder, B., & Grèzes, J. (2008). Emotional modulation of visual and motor areas by dynamic body expressions of anger. Social Neuroscience, 3, 199–212. http://dx.doi.org/10.1080/17470910701394368

Pichon, S., de Gelder, B., & Grèzes, J. (2009). Two different faces of threat. Comparing the neural systems for recognizing fear and anger in dynamic body expressions. NeuroImage, 47,1873–1883. http://dx.doi.org/10.1016/j.neuroimage.2009.03.084

Pichon, S., de Gelder, B., & Grèzes, J. (2012). Threat prompts defensive brain responses independently of attentional control. Cerebral Cortex, 22, 274–285. http://dx.doi.org/10.1093/cercor/bhr060

Ponseti, J., Bosinski, H. A., Wolff, S., Peller, M., Jansen, O., Mehdorn, H. M.,… Siebner, H. R. (2006). A functional endophenotype for sexual orientation in humans. NeuroImage, 33, 825–833. http://dx.doi.org/10.1016/j.neuroimage.2006.08.002

Ratner, S. C. (1967). “Comparative aspects of hypnosis,” in Handbook of Clinical and Experimental Hypnosis, ed J. E. Gordon (New York: Macmillan), 550–587.

Roelofs, K., Minelli, A., Mars, R. B., van Peer, J., & Toni, I. (2009). On the neural control of social emotional behavior. Social cognitive and affective neuroscience, 4(1), 50–58. http://dx.doi.org/10.1093/scan/nsn036

Rousselet, G. A., Pernet, C. R. & Wilcox, R.R (2017). Beyond differences in means: robust graphical methods to compare two groups in neuroscience. European Journal of Neuroscience, 46, 1738–1748 https://doi.org/10.1111/ejn.13610

Saraiva, A. C., Schüür, F., & Bestmann, S. (2013). Emotional valence and contextual affordances flexibly shape approach-avoidance movements. Frontiers in psychology, 4, 933. https://doi.org/10.3389/fpsyg.2013.00933

Shackman, A. J., Salomons, T. V., Slagter, H. A., Fox, A. S., Winter, J. J., & Davidson, R. J. (2011). The integration of negative affect, pain and cognitive control in the cingulate cortex. Nature reviews. Neuroscience, 12(3), 154–167. https://doi.org/10.1038/nrn2994

Siegel, J. S., Power, J. D., Dubis, J. W., Vogel, A. C., Church, J. A., Schlaggar, B. L., & Petersen, S. E. (2014). Statistical improvements in functional magnetic resonance imaging analyses produced by censoring high motion data points. Human Brain Mapping, 35, 1981–1996. https://doi.org/10.1002/hbm.22307

Simos, P. G., Kavroulakis, E., Maris, T., Papadaki, E., Boursianis, T., Kalaitzakis, G., & Savaki, H. E. (2017) Neural foundations of overt and covert actions. NeuroImage, 52, 482–496. https://doi.org/10.1016/j.neuroimage.2017.03.036

Sinke, C.B., Sorger, B., Goebel, R., & de Gelder, B. (2010). Tease or threat? Judging social interactions from bodily expressions. NeuroImage, 49, 1717–1727. https://doi.org/10.1016/j.neuroimage.2009.09.065

Smith, S. D., & Kornelsen, J. (2011). Emotion-dependent responses in spinal cord neurons: A spinal fMRI study. NeuroImage, 58 (1), 269–274. https://doi.org/10.1016/j.neuroimage.2011.06.004

Steinmetz, P. N., Cabrales, E., Wilson, M. S., Baker, C. P., Thorp, C. K., Smith, K. A., & Treiman, D. M. (2011). Neurons in the human hippocampus and amygdala respond to both low- and high-level image properties. Journal of neurophysiology, 105(6), 2874–2884. https://doi.org/10.1152/jn.00977.2010

Stan Development Team. (2016). Stan modeling language users guide and reference manual. Retrieved from http://mc-stan.org

Straube, T., Schmidt, S., Weiss, T., Mentzel, H. J., & Miltner, W. H. (2009) Dynamic activation of the anterior cingulate cortex during anticipatory anxiety. Neuroimage. 44:975–981. https://doi.org/10.1016/j.neuroimage.2008.10.022.

Talairach, J., Bancaud, J., Geier, S., Bordas-Ferrer, M., Bonis, A., Szikla, G., & Rusu, M. (1973). The cingulate gyrus and human behaviour. Electroencephalography and Clinical Neurophysiology, 34, 45–52.

Talairach, J.; & Tournoux, P. (1998). Co-planar stereotaxic atlas of the human brain. Thieme, New York.

Tomasino, B., Weiss, P. H., & Fink, G. R. (2012). Imagined tool-use in near and far space modulates the extra-striate body area. Neuropsychologia. 50, 2467–2476. https://doi.org/10.1016/j.neuropsychologia.2012.06.018

Urgesi, C., Berlucchi, G., & Aglioti, S.M. (2004). Magnetic stimulation of extrastriate body area impairs visual processing of nonfacial body parts. Current Biology, 14, 2130–2134. https://doi.org/10.1016/j.cub.2004.11.031

Van den Stock, J., Tamietto, M., Sorger, B., Pichon, S., Grézes, J., & de Gelder, B. (2011). Cortico-subcortical visual, somatosensory, and motor activations for perceiving dynamic whole-body emotional expressions with and without striate cortex (V1). Proceedings of the National Academy of Sciences of the United States of America, 108, 16188–16193. https://doi.org/10.1073/pnas.1107214108

Van den Stock, J., Hortensius, R., Sinke, C., Goebel, R., & de Gelder, B. (2015). Personality traits predict brain activation and connectivity when witnessing a violent conflict. Scientific reports, 5, 13779. https://doi.org/10.1038/srep13779

Van Loon, A. M., van den Wildenberg, W. P. M., van Stegeren, A. H., Ridderinkhof, K. R., & Hajcak, G. (2010). Emotional stimuli modulate readiness for action: A transcranial magnetic stimulation study. Cognitive, Affective, & Behavioral Neuroscience, 10, 174–181. https://doi.org/10.3758/CABN.10.2.174

Vogt, B. A., Berger, G. R., & Derbyshire, S. W. (2003). Structural and functional dichotomy of human midcingulate cortex. The European Journal of Neuroscience, 18, 3134–3144. https://doi.org/10.1111/j.1460-9568.2003.03034.x

Vogt, B. A. (2005). Pain and emotion interactions in subregions of the cingulate gyrus. Nature Reviews Neuroscience, 6, 533–544. https://doi.org/10.1038/nrn1704

Vogt, B. A., Vogt, L., & Laureys, S. (2006). Cytology and functionally correlated circuits of human posterior cingulate areas. NeuroImage, 29, 452–466. https://doi.org/10.1016/j.neuroimage.2005.07.048

Vogt, B. A., & Vogt, L. J. (2009). Opioids, placebos and descending control of pain and stress systems. In B. A. Vogt (Ed), Cingulate neurobiology and disease (pp. 339–364). London: Oxford University Press.

Vogt, B. A. (2016) Midcingulate cortex: Structure, connections, homologies, functions and diseases. Journal of Chemical Neuroanatomy, 74, 28–46. https://doi.org/10.1016/j.jchemneu.2016.01.010

Volchan, E., Souza, G. G., Franklin, C. M., Norte, C. E., Rocha-Rego, V., Oliveira, J. M., … Figueira I. (2011). Is there tonic immobility in humans? Biological evidence from victims of traumatic stress. Biological Psychology, 88 (1), 13–19. https://doi.org/10.1016/j.biopsycho.2011.06.002

Volchan, E., Rocha-Rego, V., Bastos, A. F., Oliveira, J. M., Franklin, C., Gleiser, S., … Figueira, I. (2017). Immobility reactions under threat: a contribution to human defensive cascade and PTSD. Neuroscience and Biobehavioral Reviews, 76, 29–38. https://doi.org/10.1016/j.neubiorev.2017.01.025

Zimmermann, M., Mars, R. B., de Lange, F. P., Toni, I., & Verhagen, L. (2017). Is the extrastriate body area part of the dorsal visuomotor stream? Brain structure & function, 223(1), 31–46. https://doi.org/10.1007/s00429-017-1469-0

Wilcox, R. R. (2016). Comparing dependent robust correlations. The British Journal of Mathematical Statistical Psychology, 69, 215–224. https://doi.org/10.1111/bmsp.12069.

